# Aging causes changes in transcriptional noise across a diverse set of cell types

**DOI:** 10.1101/2022.06.23.497402

**Authors:** G. Edward W. Marti, Steven Chu, Stephen R. Quake

## Abstract

Aging and its associated diseases result from complex changes in cell state which can be examined with single-cell transcriptomic approaches. We analyzed gene expression noise, a measure of cellular heterogeneity, across age and many cell types and tissues using the single cell atlas *Tabula Muris Senis*, and characterized the noise properties of most coding genes. We developed a quantitative, well-calibrated statistical model of single-cell RNAseq measurement from which we sensitively detected changes in gene expression noise. We found thousands of genes with significantly changing gene expression noise with age. Not all genes had increasing noise with age—many showed a robust decreases of noise. There were clear biological correlation between subsets of genes, with a systemic decrease of noise in oxidative phosphorylation pathways while immune pathways involved in antigen presentation saw an increase. These effects were seen robustly across cell types and tissues, impacting many organs of healthy, aging mice.

## INTRODUCTION

There has been longstanding interest in whether gene expression noise changes with age, and whether there is a causal relationship between the two. Does aging cause gene expression to become dysregulated and hence noisier? Does noisy gene expression have biological consequences which in turn cause diseases of aging? These are tantalizing questions, and unfortunately the literature contains conflicting results even on the most fundamental question of whether gene expression noise changes with age, regardless of cause and effect. Prior work in cells of the heart have indicated increases in noise for some genes [1], while other work in immune cells of the blood have shown no change in age [2]. It is of course possible and perhaps even likely that both results are correct and that changes in gene expression noise are cell type dependent.

The recent development of whole-organism cell atlases provide an unprecedented view of change in the transcriptome with age and disease [3, 4]. These atlases demonstrate that cell types and states are fluid, changing with age, disease, and lineage in complex ways[5–9]. Aging cell atlases contain critical evidence of changes in cell type specialization, clonal expansion, and heterogeneous cellular response to stress and damage [10]. Deciphering changes in the variation or continuum of gene expression is challenging because substantial quantities of technical noise and systematic errors that occur during various steps of the single-cell mRNA sequencing pipeline [11–20].

We sought to systematically and comprehensively address such questions for all expressed genes by analyzing gene expression noise as a function of age across large numbers of cell types and tissues using the single cell atlas Tabula Muris Senis. A major challenge in this work was to develop new statistical techniques which faithfully model, quantify, identify, and deconvolve measurement noise from true biological gene expression noise. We developed a quantitative, well-calibrated statistical model of single-cell RNAseq measurement from which we calibrated molecule numbers, inferred the total mRNA content of cells, and sensitively detected changes in gene expression noise in low and moderately expressed genes.

When these methods are applied to *Tabula Muris Senis*, we found thousands of genes with significantly changing gene expression noise with age. Interestingly, not all genes had increasing noise with age—some had robust decreases of noise with age. There are clear biological relationships between subsets of these genes, with energy pathways involved in oxidative phosphorylation seeing a decrease in noise while immune pathways involved in antigen presentation saw an increase in noise. These effects were seen robustly across cell types and tissues, impacting many organs of normally aging mice. In contrast to the statistically significant correlations, the biological significance of the changes in the control of gene regulation are yet to be understood.

## RESULTS

### A model of mRNA capture and amplification quantifies technical noise in single-cell RNAseq

Quantifying biological variation in gene expression from single-cell transcriptomic (scRNAseq) data requires calibrating the technical noise contribution of the overall measured noise [13]. Technical noise depends on gene expression and the total number of captured molecules, both of which change with age and cell state. To address these challenges, we developed a computational approach based on a simple model of mRNA capture and amplification to distinguish true biological sources of gene expression noise from technical noise sources (Fig. 1).

**FIG. 1:**
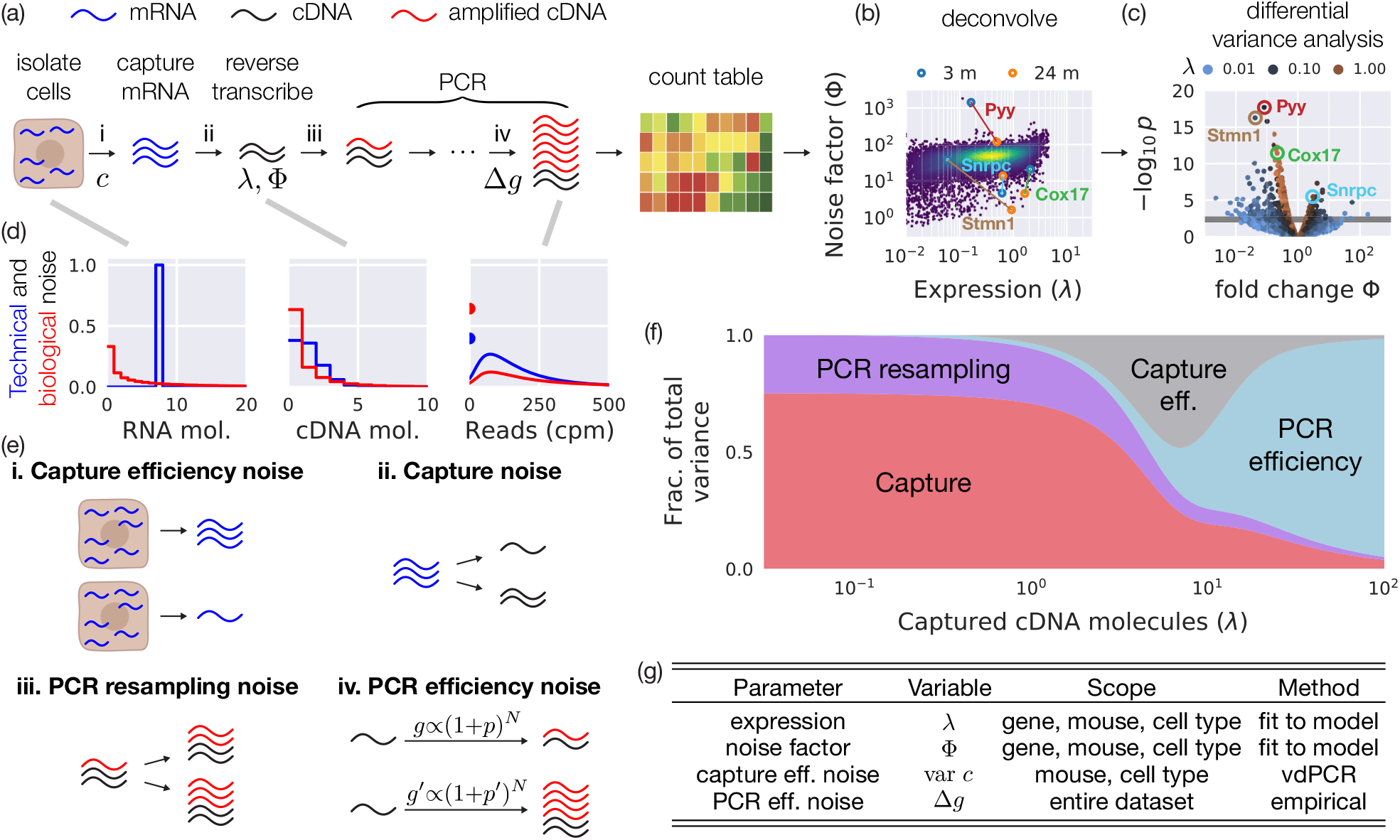
First principles statistical model single-cell mRNA sequencing. (a) A schematic of our first principles model of mRNA capture and reverse transcription and amplification. (b) Our model allows us to deconvolve expression into estimates of the mean number of cDNA molecules (*λ*) and their noise factor Φ, from which we can (c) perform differential analysis of changing gene expression noise with age. (d) Our model provides a quantitative prediction of the distribution of mRNA and cDNA molecules after each step for purely technical noise (blue) and both technical and biological noise (red). Shown here is a negative binomial distribution of mRNA, though estimates of Φ are generic and do not depend on the specific biological noise model. A mathematical formulation is included in the supplemental material. (e) Technical noise enters in four steps, as (**i**) variation in the capture efficiency, or total mRNA content, per cell, (**ii**) finite probability of capturing and reverse transcribing any given mRNA molecule, (**iii**) finite probability that a molecule is amplified during each step of PCR, and (**iv**) well-to-well variations of in PCR efficiency. (e) Quantitative contributions of variance from each technical noise term for pancreatic *β* cells (var *c* = 0.2 and Δ*g* = 0.5, see text for explanation). Capture and PCR resampling noise dominate for low expressed genes, less than five cDNA per well, while PCR efficiency noise dominates for higher expressed genes.

Our model has two key steps, (1) capture and reverse transcription to cDNA, and (2) amplification of cDNA by PCR. mRNA is captured from cells that have been isolated, sorted, and lysed from dissociated tissues of mice of several ages, from 3 m to 24 m [4]. The total accessible mRNA content per cell varies substantially between cells within the same tissue and varies systematically between cells isolated from mice of different ages. We define technical noise as the measured distribution of reads arising from a fixed and noiseless (true) number of mRNA molecules per cell, while biological noise arises from a cell-to-cell variation in mRNA content. mRNA is captured (Fig. 1e, **i**) and reverse transcribed (Fig. 1e, **ii**) stochastically. The observed number of cDNA molecules *n* for a given gene is then given by the Poisson distribution,

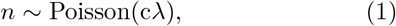

where *c* is the relative capture/conversion efficiency of RNA to cDNA, and *λ* is the mean number of cDNA molecules per cell of that gene. We estimate from exogenous spike-in RNA that at most 18% of starting mRNA transcripts are reverse transcribed and then detected (Fig. S3).

PCR amplification is critical to detect small numbers of cDNA molecules but adds additional technical noise and distortion. cDNA molecules are resampled and amplified through many cycles of PCR, which adds PCR resampling noise (Fig. 1e, **iii**). In addition, PCR efficiency may vary slightly from well to well due to slight differences in reagents, temperature, evaporation rates, etc., which lead to a long-tailed distribution in the number of amplified molecules (Fig. 1e, **iv**). These processes are quantitatively modeled with a single parameter representing variation in PCR efficiency: Δ*g* = 0.5, which was empirically estimated for the entire dataset (Methods and Supplemental Material). Finally, we assume the library preparation, sequencing, alignment, and normalization to counts per million (cpm) adds minimal additional noise compared to the previous steps (Supplemental Material Sec. I A).

A full quantitative model is described in the Methods. In brief, the counts per million *x* for a given gene depends on the relative capture efficiency *c*, the PCR gain *g*, the stochastic PCR noise *y*, and a noise factor Φ (Methods and Supplemental Material for details).

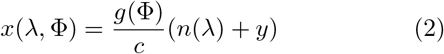

The distribution *P*(*x* |*λ*, Φ) can be solved by integrating over all distributions. We then fit the measured mean and variance of log_10_(1 + cpm) to the theoretical mean and variance of log_10_(1 + *x*(*λ*, Φ)), from which we determine estimates of *λ* and Φ for each gene.

Despite the distortion of PCR, the dominant source of technical noise for most genes is the random sampling of molecules (Fig. 1f). Typically 80-97% of genes are expressed with mean expression of less than one cDNA molecules per well (Fig. S5). A protocol that adds unique molecular identifiers (UMIs) at the expense of capture efficiency, typically a threefold change in efficiency in most protocols [21], will negatively impact the estimation of the noise factor Φ for low expressed genes (Supplemental Material). Only at high expression levels, greater than five cDNA molecules per well, does PCR efficiency noise begin to dominate. A protocol that uses UMIs to remove PCR efficiency noise would improve estimates of Φ for these highly expressed genes, as has been used in previous work [13, 14]; however only a small fraction of the genes are sufficiently highly expressed to benefit (Fig. S5).

For any given cell type and gene the model has excellent quantitative agreement with the data with only four fit parameters: expression and noise level for the gene in the cell type, a variance of cell sizes (total mRNA content) for each cell type and mouse, and a single global parameter describing the long-tailed distribution of the molecule number after PCR amplification (Fig. 1g).

### Low expression, high dropout genes and exogenous spike-in controls validate and calibrate the model

We validated the technical noise model through three computational approaches that dissect different stages of the single-cell pipeline (Fig. 2a). First, we performed virtual digital PCR (vdPCR) on the measured data, a computational approach analogous to digital PCR [22]. In vdPCR, the count table is binarized, with entries equal to one when reads for a gene are present, and zero otherwise (Fig. 2b and Supplemental Material Sec. I A). Second, we estimate the mean expression of a single cDNA molecule and the total number of cDNA molecules and mRNA in each sample. Finally, we estimate the long-tailed distribution of reads after PCR amplification from each cDNA molecule, a key source of measurement noise. All three approaches are applied to high dropout, low expression genes as well as exogenous spike-in RNA (ERCCs after the External RNA Controls Consortium, from which they were developed).

**FIG. 2:**
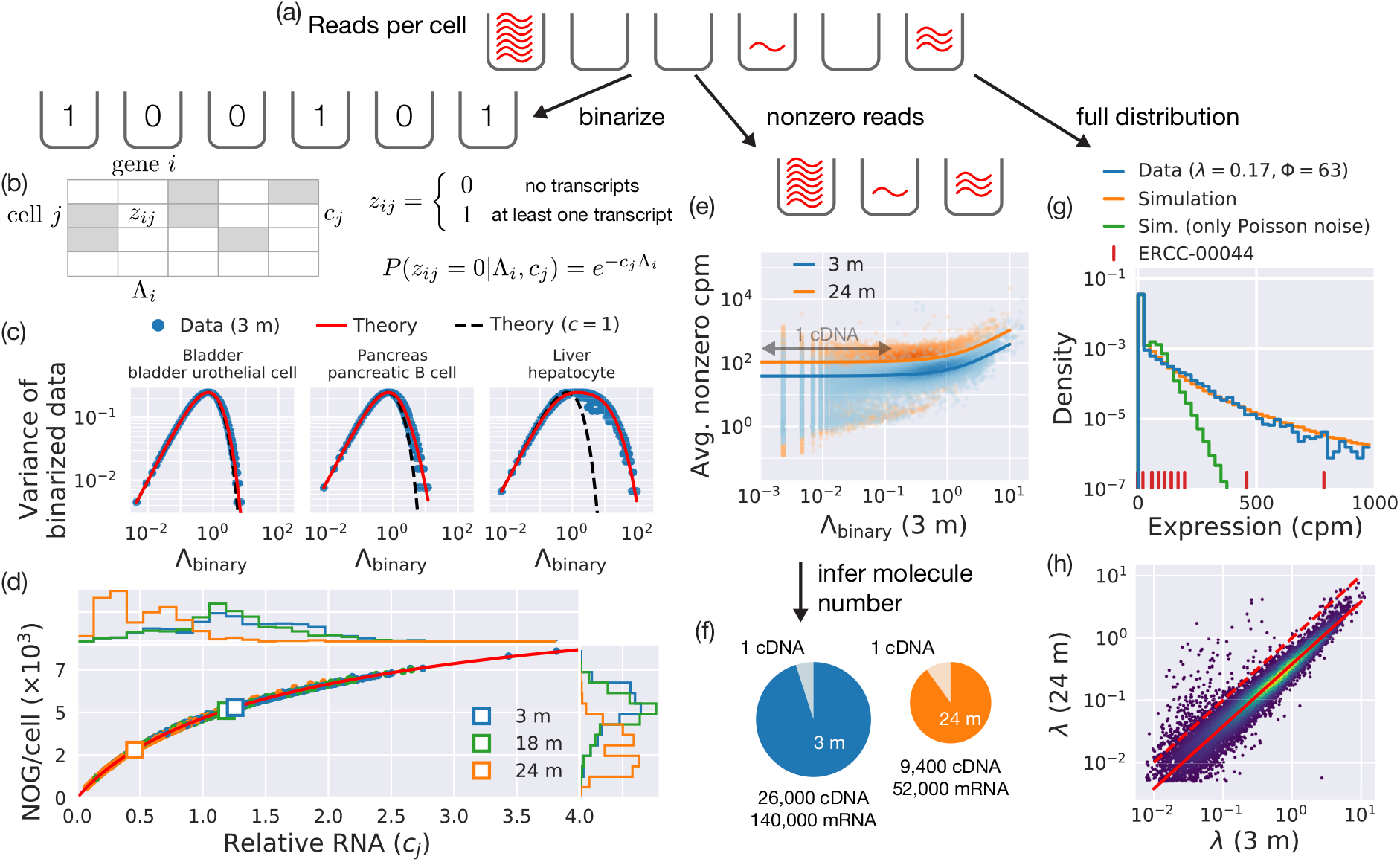
Validation and calibration of statistical model. (a) The statistical model is calibrated by the large number of low-expressed, high dropout genes present in RNAseq data. Reads per cell for a given gene are represented as proportional to quantities of amplified DNA reads in different wells for one cell type. (b) Reads are binarized so any aligned transcripts (nonzero reads) are represented as a one, and dropouts as a zero. These data are sensitive to noise accrued during mRNA capture and reverse transcription, but not PCR distortion. (c) Capture efficiency noise leads to an increase in the variance of binarized data (blue points for each gene), shown here in the increasingly heterogeneous populations of urothelial cells, pancreatic *β* cells, and hepatocytes. The data is well described by a single parameter (solid red curve), var *c*, and disagrees with a model that assumes all cells have the same capture efficiency (dashed black line) (d) The number of unique genes (NOG) per cell is entirely explained by relative capture efficiency, equivalent to the relative RNA content, with a decrease in NOG with age predicted by their relative decrease in captured mRNA content per cell (solid red line). Histograms of relative capture efficiency (cell size) and NOG are show on the top and right, respectively. Each point is one pancreatic *β* cells. Cells isolated from 3 m, 18 m, and 24 m mice are blue, green, and orange, respectively, with squares representing the population mean. (e) Low expressed genes with high dropout are a well-calibrated source of single molecules. The few wells that contain transcripts most likely contain only one molecule. Averaging nonzero reads of these genes estimates the counts per million (cpm) associated with one cDNA molecule, here shown for pancreatic *β* cells. The mean cpm of nonzero reads is largely independent of expression for Λ *<* 0.3. Cells from 24 m mice show a higher cpm per molecule than cells from 3 m mice because we reverse transcribe less cDNA, and each cDNA is correspondingly a larger fraction of the limited number of molecules. (f) We estimate 2.6 *×* 10^4^ and 9.4 *×* 10^3^ reverse transcribed molecules of cDNA, corresponding to 1.4 *×* 10^5^ and 5.2 *×* 10^4^ molecules of accessible mRNA, isolated from the from pancreatic *β* cells of 3 m and 24 m mice, respectively. (g) The distribution of expression for genes with low expression (*λ ≈* 0.17) and similar noise factor (Φ *≈* 63) is long-tailed, and is fit well with a model including PCR efficiency noise (Fig. 1e, **iv**). The distribution is validated by exogenous RNA spike-ins (ERCCs), which show a consistent long-tailed distribution (red ticks). (h) Estimates of expression in 24 mice are systematically smaller than for 3 m mice across all measured gene expressions. This effect is included and removed during differential noise analysis.

The vdPCR count table includes noise from the cell-tocell size variation (capture efficiency noise) and stochastic capture (Fig 1e **i** and **ii**) but not PCR resampling or amplification. Dropouts, in which a gene is not detected in a given cell, is due to the Poisson statistics of capture. It has previously been shown that dropouts are unlikely to distort the true distribution of gene expression. In other words, the number of zeros found in single cell sampling is not inflated beyond what is expected by Poisson sampling [23].

We find that a model with a single parameter describing capture efficiency explains the variance of vdPCR data for the vast majority of genes in a given cell type (Fig. 2c) (Supplemental Material Sec. I A). This parameter can also be interpreted as representing that different cells have different quantities of total mRNA. The variance of vdPCR data increases and then decreases with estimated gene expression, as highly expressed genes have nearly no dropouts and therefore vanishing variance in vdPCR (Fig. 2c, blue points). Compared to the simple model without noise in capture efficiency, the data shows a shift in the curve at high expressions due to such noise (Fig. 2c, red line and dashed black line), and the actual data is well fit with a single parameter describing capture efficiency noise.

This parameter varies with cell type and mouse, from 0.07 for urothelial cells, 0.18 for pancreatic *β* cells, and 1.2 for hepatocytes (Fig. 2c, measured for the 3 m mouse with the most cells). We have also derived similar values for this parameter from other data sets by a reanalysis of previous literature. That analysis led to values of 0.12 for U2OS cells in merFISH expansion microscopy [24] and 0.13 for mouse embryonic stem cells in CEL-seq [13] (Supplemental Material Sec. I C). Whether the changing heterogeneity shown here is a purely technical effect or demonstrates true biological noise is unclear. Regardless, we treat it as technical noise in this analysis.

The relative capture efficiency (total mRNA content) and number of unique genes (NOG) from cells isolated from 3 m and 18 m agree well (Fig. 2d, blue and green), while cells from 24 m mice show a substantial decrease in both metrics (orange). Our model shows excellent quantitative agreement of the observed relationship between relative mRNA content and NOG (solid red line). The decreased NOG seen in cells isolated from 24 m mice has been observed before [4], but our analysis demonstrates that it is due entirely to the decreased number of captured mRNA molecules in those cells. While the origins of this systematic decrease in cDNA are unknown, the shift is consistent across three orders of magnitude of gene expression (Fig. S4), suggesting that mRNA from older tissues is likely more degraded during tissue handling and dissociation. However, this interpretation is not definitive and could have a biological origin as well, for instance due to smaller or less transcriptionally active cells in older mice. Regardless of origin, in this work we consider the decrease in mRNA and cDNA to be a systematic error that must be accounted for to explore true variations in noise with age.

The distribution of reads from the large number of high dropout genes gives a detailed picture of the PCR amplification process and its technical noise. Measured reads from a low expressed gene with high dropout most likely originate from a single cDNA molecule. We test this assumption by averaging the mean expression of only cells with nonzero reads (Fig. 2e, each point is one gene). The result is independent of expression for 10^−3^ *<* Λ *<* 10^−1^, here shown for pancreatic *β* cells, demonstrating that in this range of data only one cDNA molecule is captured and amplified (Λ is the expression estimated from dropout, see Supplemental Material). This further confirms that dropouts are reflective of the Poisson statistics of molecule capture and are not zero inflated [23]. The average cpm per molecule is significantly larger for 24 m mice because fewer total cDNA molecules are captured (Fig. 2f). The geometric mean from one molecule is 50 cpm in 3 m mice and 110 cpm in 24 m mice, suggesting that there the number of accessible mRNA molecules per pancreatic beta cell decreases from 110,000 and 50,000 as the age increases from 3 m to 24 m, assuming a capture efficiency is 18%.

The distribution of reads is long-tailed because of cell-to-cell variations in the PCR efficiency (Fig. 2g). A model of stochastic capture and sampling during PCR alone (Fig. 1b **iii**) does not predict the long tails seen in moderately expressed genes (blue curve, 731 genes of similar expression and noise factor) and in spike-in, exogenous RNA (ERCC-00044, red vertical ticks). We model the PCR gain per cell (cpm per cDNA molecule) as a log-normal distribution with a single parameter Δ*g* = 0.5 − 0.8, corresponding to a 6-10% variation of PCR efficiency per cycle, and observe excellent quantitative agreement (orange line).

Our model explains the distributions and noise of the majority of low expression genes using a small number of fit parameters (Fig. 2). We calculate a capture efficiency noise term var *c* for each cell type and mouse from vd-PCR data, and empirically use Δ*g* = 0.5 to model PCR efficiency noise for the entire dataset. The remaining two parameters are the expression *λ* (the mean number of cDNA molecules) and the noise factor Φ of each gene, in each mouse, for each cell type, fit by matching the summary statistics (mean and variance of log_10_(1 + cpm)) of the observed data to expected values from the quantitative model (Supplemental Material Sec. I D). Fitting data to *λ* and Φ, which are uncorrelated, deconvolves changes in expression with changes in technical noise (Fig. 1a and Fig. S2). This approach is similar in spirit to variance stabilizing transforms that decorrelate noise with expression [25–27], but our method works equally well for low (*λ <* 1) or high (*λ >* 1) expression.

The noise factor Φ, an estimate for the measured noise in cDNA expression for each gene, is equal to the product of the unknown mean PCR gain *G*_PCR_ (cpm per cDNA molecule) and a biological noise term *η*_biol_ that depends on the distribution of true biological noise, Φ = *G*_PCR_*η*_biol_. Two limiting cases of bursting gene expression [28–30] are the zero inflated Poisson distribution of mRNA, for which *η*_biol_ is the Fano factor (variance over mean), and the negative binomial distribution, for which *η*_biol_ is approximately the square root of the Fano factor (Supplemental Material). Since the PCR gain is unknown but is expected not to vary with age, we calculate the fold change of Φ between cells isolated from mice of different ages to remove its effect (Fig. 1c).

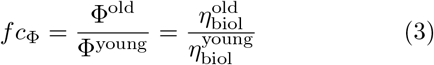

Uncertainty in *fc*_Φ_ is estimated from the technical noise model and from the biological replicates of mice of the same age (Fig. S1). The quantitative contribution of different noise sources to technical gene expression noise are shown in Fig. 1f. For high expression genes, PCR efficiency dominates and estimating technical noise becomes increasingly imprecise. Similarly, low expression genes with very high dropout have insufficient statistical power to determine Φ. For this reason, our analysis excludes genes with *λ >* 3 and *λ <* 10^−2^.

### Differential noise genes (DNGs) are enriched in pathways relating to oxidative phosphorylation and antigen processing and presentation

Our model-based analysis is applied to each gene, for each cell type, and for each mouse across in the TMS data set[4]. TMS-defined cell types are sub-clustered for a more homogeneous population and filtered for those with at least two 3 m mice and two 24 m mice and at least 150 cells mouse (Fig. S1). Batch effects are minimized by performing estimation of Φ within individual mice, normalizing expression and noise by the relative expression per mouse to account for changes in total cDNA, and aggregating by age (Fig. S1). A summary of results is show in Fig. 3.

**FIG. 3:**
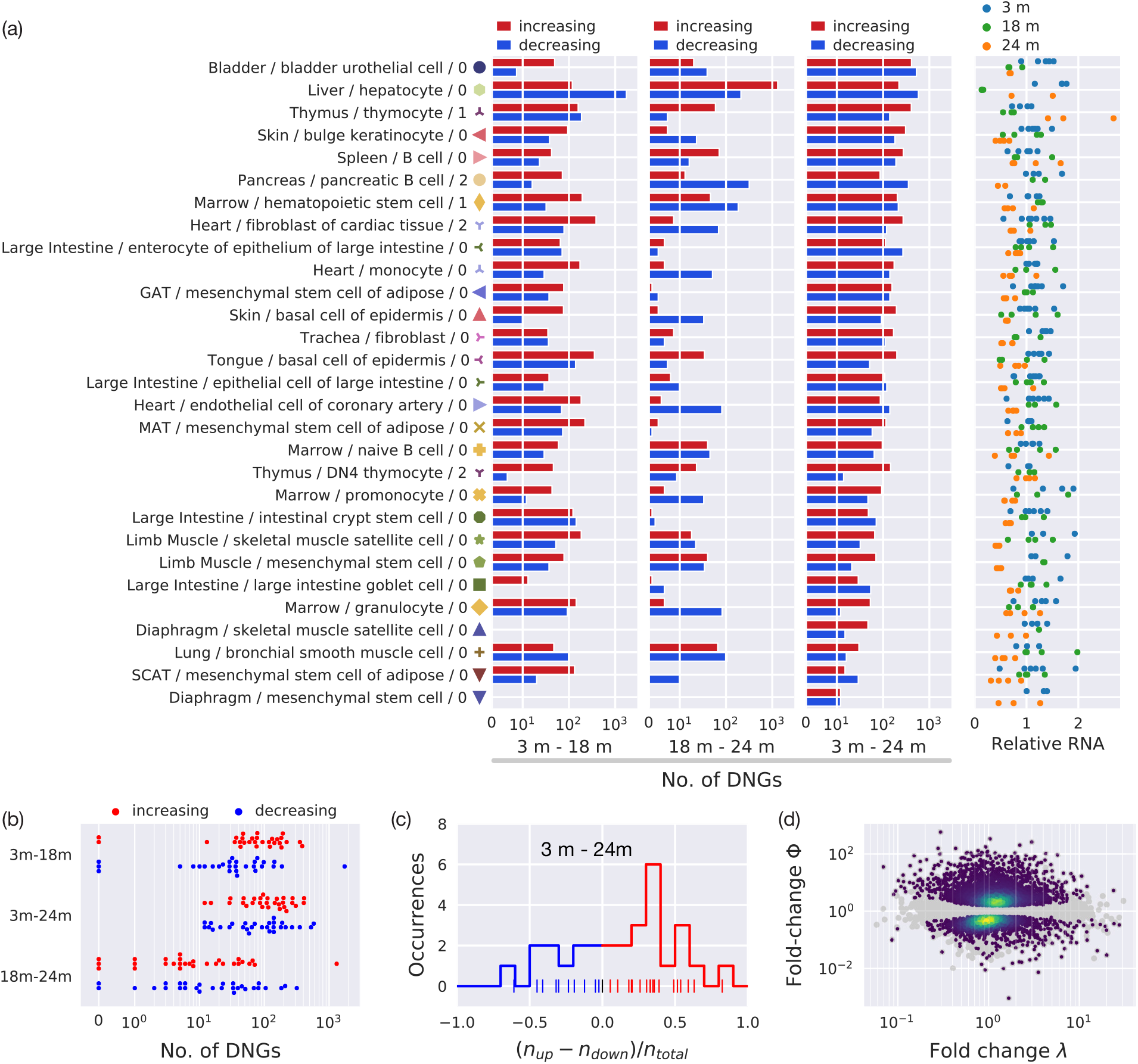
Number of DNGs across all tissues. (a) 29 cell types in 17 tissues pass our quality control of at least two mice per age and at least 150 cells per age. Typically 10-300 DNGs and 50-2000 DEGs per cell type between cells isolated from 3 m and 24 m mice. Cells isolated from 24 m mice show a systematic decrease in relative mRNA content. (b) A large number of DNGs are found between in the 3 m-18 m and 3 m-24 m comparison, with many fewer between 18 m-24 m. (c) Most cell types have more DNGs corresponding to increasing variation or noise than decreasing. (d) Fold change of gene expression (*λ*) and noise factor (Φ) between cells isolated from 3 m and 24 m mice, including all cell types and tissues. The colored dots denote the density of (statistically significant genes), as compared to all genes (gray and colored dots).

We observe substantially more differential noise genes (DNGs), genes that shown significant fold change in the noise factor, between 3 m and 24 m, and the fewest number of DNGs between 18 m and 24 m (Fig. 3b). These data suggest a marked and systemic change in biological gene expression noise or biological variation between 3 m and 18 m, with more organ-or cell-specific changes towards 24 m. Liver hepatocytes stand out with particularly large number of DNGs between 18 m and 24 m. This analysis find typically 50% more DNGs corresponding to increasing than decreasing noise (Fig. 4c). DNGs are corrected for decreasing relative mRNA in older mice (Fig. 4b).

**FIG. 4:**
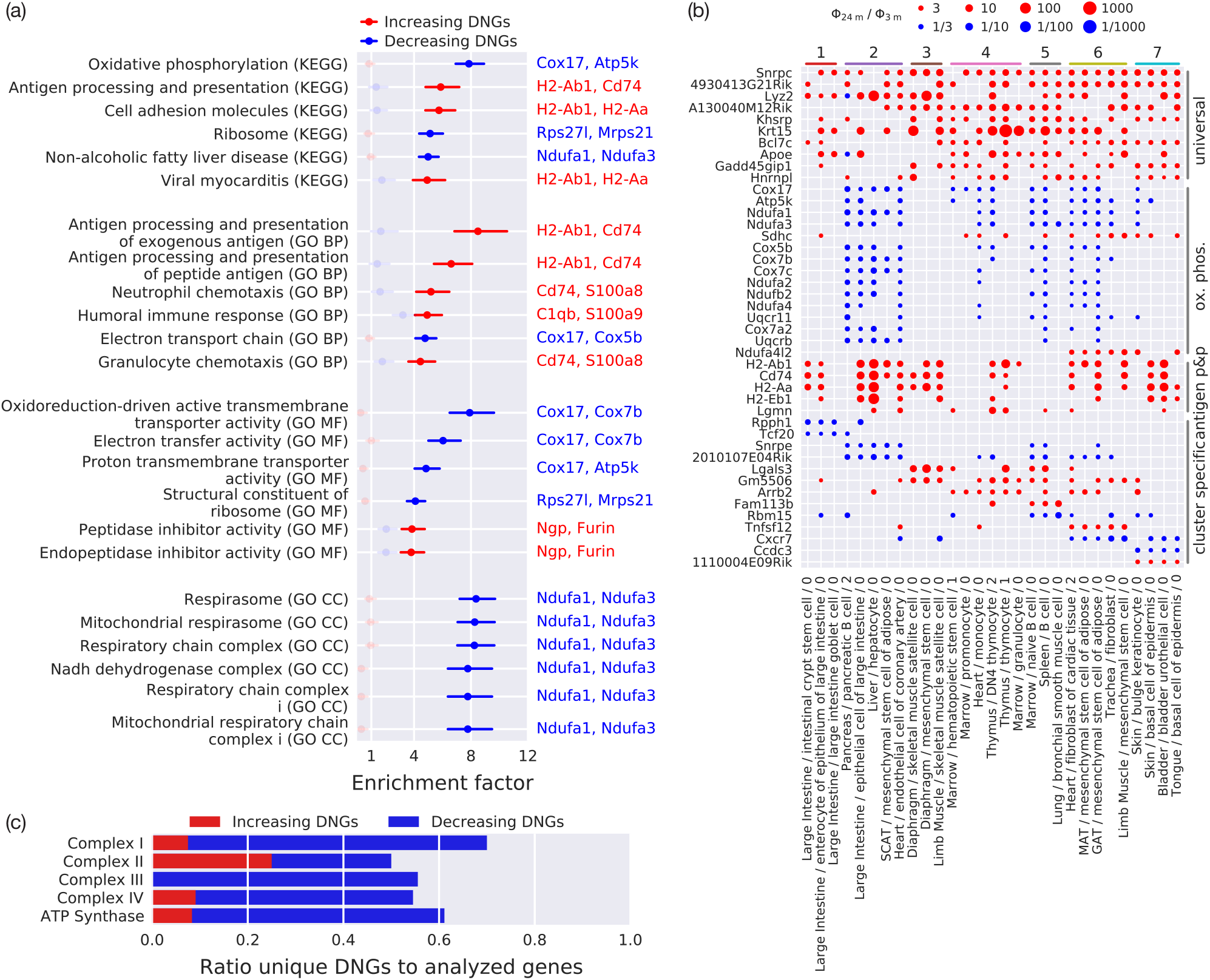
Pathway and gene ontology enrichment of increasing and decreasing DNGs. (a) Pathways pertaining to oxidative phosphorylation and antigen processing and presentation are the most frequently enriched in decreasing and increasing DNGs, respectively. Other pathways (virial myocarditis, non-alcoholic fatty liver diseases) appear because they overlap with oxidative phosphorylation or antigen processing and presentation, as can be seen by the most frequently seen genes in the pathway are indicated on the right (increasing DNG in red, decreasing in blue). The top enriched pathways are significantly enriched in either increasing or decreasing DNGs, but never both. Enrichment scores are calculated based on the frequency of DNGs found across tissues and cell types as compared to the frequency of analyzed genes for which Φ can be accurately determined (0.05 *< λ <* 3), with 1 corresponding to no enrichment. (b) Occurrence of DNGs in each cell type. Genes are grouped as universal (genes found in more than 25% of cell types), oxidative phosphorylation (oxphos), antigen processing and presentation (antigen p&p), and DNGs specific to each cluster. Increasing DNGs are red, decreasing DNGs are blue. (c) Ratio of DNGs to analyzed genes with calculated Φ for each major component of the electron transport chain. The majority of analyzed genes are found as DNGs, indicating a widespread change in the regulation of oxidative phosphorylation. While most DNGs are decreasing, complex II contains several increasing DNGs, notably *Sdhc*.

Cell types may share up to 50% of the same DNGs (Fig. 4e). We cluster based on DNG similarity (using the Jaccard index) and observe clusters with loosely similar function or origin, including crypt cells in cluster 1, and immune cells in clusters 4 and 5, mesenchymal stem cells in cluster 6, and epithelial cells in cluster 7. The number of DNGs is weakly correlated with the number of DEGs. Cell types with more DEGs and DNGs do not show an increase in the number of overlapping genes that are both DEGs and DNGs, with a Jaccard similarity of typically 3-13% (Fig. S7).

A set of “universally varying” DNGs appear frequently across the 29 cell types and 17 tissues that pass our quality control, with 129 genes appearing in more than 8 (28%) of the cell types (the family-wise error rate (*FWER*) of making a false discovery is ≪ 0.01).) An enrichment analysis of DNGs against the KEGG and GO gene databases shows a significant increase in enrichment in genes relating to oxidative phosphorylation and antigen processing and presentation (Fig. 4a, *p* ≪ 10^−3^, see Supplemental Material). Examples of “universal” DNGs, DNGs enriched in oxidative phosphorylation and antigen processing and presentation, and genes specific to clusters of cell types are shown in Fig 4b. Nearly all of these genes show a consistent increase or decrease in their noise factor across the cell types. DNGs related to antigen processing and presentation tend to have increasing noise factor with age, while nearly all DNGs involved in oxidative phosphorylation show a counter-intuitive *decrease* of noise with age.

Not only is oxidative phosphorylation enriched in DNGs, but the majority of analyzed genes in all major components of the electron transport chain are DNGs (Fig 4c). *Ndufa1* in Complex I (NADH ubiquinone oxidoreductase), *Cox17* in Complex IV (cytochrome c oxidase), and *Atp5k* in ATP synthase are decreasing DNGs found in half of the cell types studied here. These genes typically show a significant decrease in Φ between 3 m and 18 m, with either a continued decrease in Φ towards 24 m (in cluster 2) or a flattening between 18 m and 24 m (cluster 4), depending on cell type (Ext. Fig. S8). This is a striking result since mitochondrial dysregulation is a well known hallmark of aging.

“Inflammaging”, chronic inflammation that correlates with aging and many age-related diseases, is seen in nearly all single-cell mRNA seq studies of aging [cite]. We identify many related pathways, most notably antigen processing and presentation, as having genes with increased noise (Fig. 4a). The most frequently found DNGs in these pathways are related to MHC class II presentation, including *Cd74, Lgmn*, and the antigen binding genes *H2-Aa* (orthologous to *HLA-DQA1* in humans), *H2-Ab1* (*HLA-DQB2*), and *H2-Eb1* (*HLA-DRB1*). We observe increasing DNGs (increasing biological variation) across a wide range of cell types, including epithelial cells of the large intestine and bladder, endothelial cells in the heart, hepatocytes, thmyocytes, and mesenchymal stem cells across many tissues. Several of these non-canonical presenters of MHC class II have been observed during inflammation, aging, or stress, including in hepatocytes [31, 32], thymocytes [33], epithelial cells [34], and endothelial cells [35]. Canonical presenters of MHC class II molecules, professional antigen producing cells such as B cells, macropaghes, and dendritic cells, are excluded from differential variance analysis for these genes because their expression of MHC class II molecules is far too high to accurately determine Φ (*λ* ≫ 3).

### Decrease variation of oxidative phosphorylation is correlated with well-known transcription factors and regulators

We hypothesise that the broad decrease of variation in oxidative phosphorylation across cell types and tissues may be driven by a few regulator genes and transcription factors. We therefore developed a noise score for genes in the oxidative phosphorylation pathway based on a weighted sum of relevant genes, where a larger score corresponds to decreasing noise (Fig. 5a). We then search for genes who changing expression across all cell types correlates the most strongly with oxidative phosphorylation noise scores (Fig. 5b). Of the twenty most correlated genes, over half are transcription and translation factors, growth factors, and regulators. A subset is show in Fig. 5c, including *Tfam* (transcription factor), *Eif2s2* (translation initiation factor), *Fibp* (growth factor), *Gps2, Dpy30*, and *Hcfc1r1* (regulator). Surprisingly, all strongly correlated genes show a decrease in expression for decreasing oxidative phosphorylation noise. A similar analysis for the antigen presentation and processing pathway is shown in Fig. S9).

**FIG. 5:**
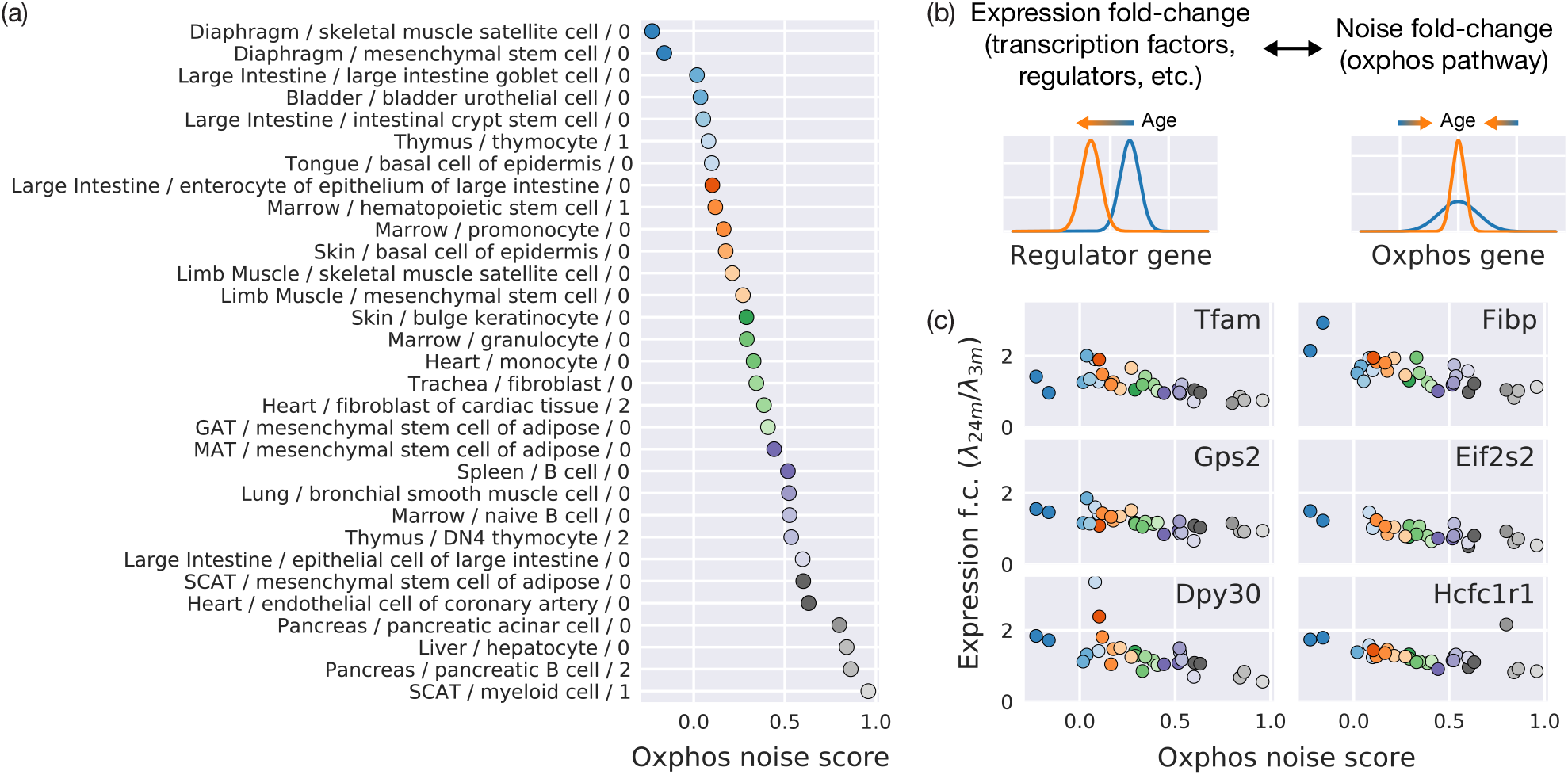
Decrease in oxidative phosphorylation variation is correlated with transcription factors and regulators. (a) Cell types in each tissue are scored (oxphos noise score) by the decrease in Φ of the most significant oxidative phosphorylation DNGs. Higher score means more decrease variation. (b) To search for drivers of oxidative phosphorylation variation, we search for genes with expression fold changes with age that correlate with the oxphos noise score. (c) More than half of the 20 most significant genes are transcriptional factors (*Tfam*), translation initiation factors (*Eif2s2*), growth factors (*Eif2s2*), or regulators (*Gps2, Dpy30*, and *Hcfc1r1*).

While it is difficult to establish causality from these correlations, these genes can now be recognized as important potential targets to decipher the mechanism of decreasing variation observed here.

## DISCUSSION

We have demonstrated a new, model-based approach to decipher biological variation from technical noise in single-cell mRNA sequencing data. Applying our method to the PlateSeq *Tabula Muris Senis* cell atlas, we observe organism-wide changes in the variation of gene expression across over key differential variance genes, genes with significant changes in gene expression variance with age in a single cell type. Hundreds of DNGs occur is over half of cell types that passed our quality control, with sufficient technical replicates (cells per mouse) and biological replicates (mice) for significance analysis. Several pathways are enriched in either increasing DNGs, corresponding to a broad-scale increase in biological variation, or decreasing DNGs that represent a homogenization of gene expression. Key pathways are those involves in oxidative phosphorylation and the electron transport chain, with a majority of decreasing DNGs, and antigen processing and presentation, with a majority of increasing DNGs.

Several types of mechanisms may drive biological variation in single-cell mRNA measurements. One possibility is that all cells of a specific cell type express the same distribution of mRNA molecules, and our measurements serve as a snapshot in time of that distribution. In this case, a DNG would indicate a change in the width of the cellular mRNA distribution. An example of this phenomenon is transcriptional noise, in which changes in gene regulation (or dysregulation) or external factors could cause a change in expression noise [cite]. Another likely possibility is that DNGs sensitively capture changes in cell state heterogeneity with age, either increasing heterogeneity (increasing DNGs) or decreasing heterogeneity (decreasing DNGs). In this view, each cell type is composed of a continuum of cell states that change with age. Many aging phenomena, including accumulated somatic DNA damage, clonal expansion, and external factors could drive cell types through multiple trajectories that is measured as a change of gene expression with age. While it is difficult to distinguish between these scenarios from single-cell RNA atlases, future work may enable observation of changes in variation due to, for instance, clonal expansion, mutant stem cells, or locally regulated factors.

Mitochondrial dysfunction is one of the hallmarks of aging [10, 36, 37]. Our observation of decreasing variation in nuclear-encoded oxidative phosphorylation genes suggests that many cells and tissues become metabolically more similar with age. Whether the loss of transcriptional heterogeneity is a cellular response to alleviate mitochondrial dysfunction, such as reduced energy output or deleterious mutations of mitochondrial DNA, or whether it is a driver that compromises the nuanced response of cells to increased stress and inflammation, remains unclear and warrants further work.

As DNGs may highlight changes in cell state heterogeneity, we postulate that inflammaging causes MHC class II antigen presentation in non-canonical MHC class II presenters, for instance hepatocytes, various epithelial cells, and thymocytes. (Fig. S9). This work suggests that such presentation is systemic and highly heterogeneous, arising to different degrees in subpopulations and thus appears as an increase biological noise. The degree to which other signs of inflammaging are present should be investigated in future work.

We acknowledge helpful discussions with E. Jerison, I. Cvijović, C. S. Peng, M. Käller, A. Isakova, A. Pisco, G. Mahmoudabadi, M. Moufarrej, M. Swift, Y. Xue, A. White, F. Zanini, U. Alon, and R. Phillips. This work was support by the Chan Zuckerberg Biohub.

## METHODS

### A minimal statistical model of scRNAseq

We evaluate a statistical model of technical gene expression noise for each cell type and mouse. To model technical noise, we assume that each cell (or well) starts with a fixed number (or narrow distribution) of *n*_*mRNA*_ molecules of a given gene. After mRNA capture and reverse transcription, there remain *n* molecules of cDNA. Each well has measurable variation in the efficiency of mRNA capture or reverse transcription, *n* = *p n*_mRNA_, which effects the total number of cDNA molecules across all genes. We define a relative capture efficiency (or cell size) *c* ∝ *p* to each cell, where ⟨*c*⟩ = 1. For convenience, we assume *c* is distribution as a log-normal distribution, though other choices of distributions yield similar results.

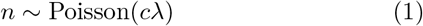

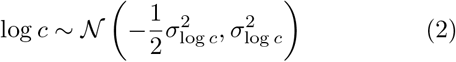

Here, 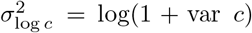, which ensures that such that *c* has a mean of 1 and variance var *c*. These terms describe technical noise that may arise from mRNA capture and reverse transcription. Because *n* depends on the product *cλ*, its Fano factor (averaged over the distribution *c* of all cells for one *λ*) is *F* = 1 + *λ* var *c*. This term can be approximated as one for small *λ*.

PCR is essential to generating enough molecules for sequencing but may distort the data. After PCR, each gene has a PCR gain *G* equal to the counts per million per cDNA molecule. Each cycle of PCR has finite efficiency so it is a stochastic process (each molecule may or may not be duplicated) and the final number of molecules is broadened compared to the input. This can be modeled as an effective input noise *y* (see Supplemental Material Sec. I B). In addition, each well may have a different efficiency *g*. Even small differences in PCR efficiency leads to substantial differences in the final molecule number after many cycles. We model *g* as having a geometric mean *G* and width Δ*g*. Experimental evidence shows this distribution is log-normal, as validated both by low expression genes and PCR.

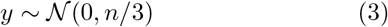

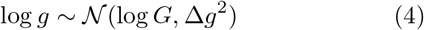

Finally, we measure a normalized counts per million *x*. By convention, *x* summed over all genes in a given cell is 10^6^.

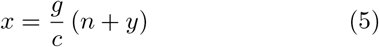

When extending the model to biological as well as technical noise, we replace *G* with Φ (see below). While the model may at first appear complicated, it has a minimum of fit parameters to prevent overfitting. We set Δ*g* = 0.5 for the entire dataset. Each mouse and cell type is described by a single parameter var *c* for all genes, describing the heterogeneity in cell sizes or capture efficiencies. Each gene, in each mouse and cell type, has only two parameters: a mean number of cDNA molecules per cell *λ* and a noise factor Φ. These few parameters capture most qualitative and quantitative aspects of gene expression noise across the entirety of *Tabula Muris Senis*.

### Virtual digital PCR

We calculate a binarized count table *z*_*ij*_ for gene *i* and cell *j* whose entries are one if there is at least one read for a given gene, and zero otherwise. The probability that *z*_*ij*_ = 0 is 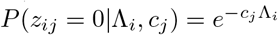, where Λ is the number of cDNA molecules (here Λ will be the vdPCR estimate of expression to be distinguished from *λ* in the full model). We use maximum likelihood estimates to calculate 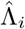 and *ĉ*_*j*_ that maximize log *ℒ* = Σ_*ij*_ *P*(*z*_*ij*_|Λ_*i*_, *c*_*j*_), where the number of observations *n*_genes_ *× n*_cells_ greatly exceeds the number of parameters *n*_genes_ + *n*_genes_ − 1 (see Supplemental Material for more details). From the estimates *ĉ*_*j*_ we can calculate the fraction of nonzero entires 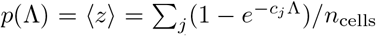 for an arbitrary gene with expression Λ, as well as the binarized variance per cell *NOG* 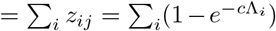 is well predicted *p*(1 *− p*) (Fig. 2c). We find that number of unique genes by the relative capture efficiency *c*_*j*_ (Fig. 2d, red line).

### Numerically evaluating the model and estimating parameters

The full distribution *P*(*x* | *λ*, Φ, var *c*, Δ*g*) of Eq. 5 may be numerically computed by integrating over *c, g, y*, and *n* (see Supplemental Material). From this distribution we calculate the theoretical mean *m*(*λ*, Φ, var *c*, Δ*g*) and variance *v*(*λ*, Φ, var *c*, Δ*g*)of log_10_(1 + *x*). We then solve for 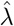 and 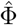 of each gene (in each mouse and cell type) such that the theoretical mean and variance match the experiment data (Extended Fig. S1b,c), as well as the technical uncertainties 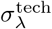 and 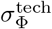 (Extended Fig. S1d). We computationally checked that estimating *λ* and Φ through the summary statistics of log_10_(1+*x*) provides much better estimates than the summary statistics of *x* because *x* has a long-tailed (log-normal) distribution. Details are given in the Supplemental Material.

### Biological gene expression noise

A gene with biological as well as technical gene expression noise will have a more complex and possibly unknown distribution of *n* that cannot be easily determined from *x*. Instead, we define a noise factor Φ = *G* = *G*_PCR_*η*_biol_, where *G*_PCR_ is the true but unknown PCR gain, and *η*_biol_ describes biological noise (see Supplemental Material for more information). We find that for several models of biological gene expression noise, our estimates of *λ* and Φ efficiently deconvolve changes in noise from changes in gene expression (see Suppelemental Material). Differential noise estimates between ages remove *G*_PCR_ and allow for comparisons of *η*_biol_ with age (Eq. 3).

### Aggregating estimates from different mice

For each gene and cell type, we make an estimate of 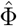 in each mouse. We aggregate estimates from all mice of the same age by taking the mean of log 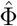 weighted by the technical estimates *σ*_log Φ_ = *σ*_Φ_*/*Φ = *CV* (Φ), where *CV* refers to the coefficient of variation. Across many orders of magnitude of gene expression, and all cell types, we observe that the measured variance 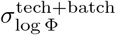 of log 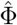 between mice of the same age is typically 2-3 times greater than 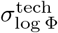, the variation expected from technical noise alone (Extended Fig. S1d). Extensive simulations show that this effect does not arise from the model or data processing (Extended Fig. S1d, second row). We use the experimentally measured distribution 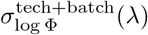 from genes of similar expression as an estimate of the confidence of Φ.

### Differential noise calculation

Aggregating log 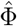 from different mice of the same age gives a noise factor and an uncertainty 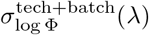. We correct for systematic changes in cDNA molecule number by normalizing the geometric mean, e.g.,

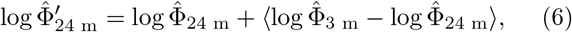

where the mean ⟨·⟩ is taken over all genes. Only genes with 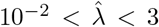 are included in the analysis. Outside of this range, the uncertainty 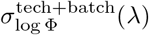 becomes large and itself uncertain.

We perform a z-test on the log fold change for each gene,

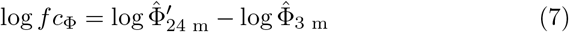

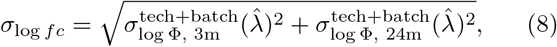

and similar for different pairs of ages. Since 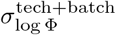 are estimated from a large number of genes of similar expression, a z-test is more appropriate than a t-test. Finally, all tests are corrected by Benjamini-Hochberg (Extended Fig. S1e). Genes that are statistically significant are termed differential noise genes (DNGs). Extensive simulations are performed on the entire pipeline to ensure results are unbiased and *p*-values are well calibrated (Fig. S1b-e, second row).

### Enrichment analysis

Many DNGs occur across multiple cell types or tissues, which we wish to weight more heavily in our enrichment factor. We calculate an enrichment score for each pathway by summing the number of DNGs in that pathway over all tissues and cell types, allowing for duplicates when a DNG is present in multiple cell types or tissues. The enrichment score is normalized by resampling all genes from which a log *fc* can be confidently calculated. More information is available in the supplemental material.

## Supplemental Figures

**FIG. S1:**
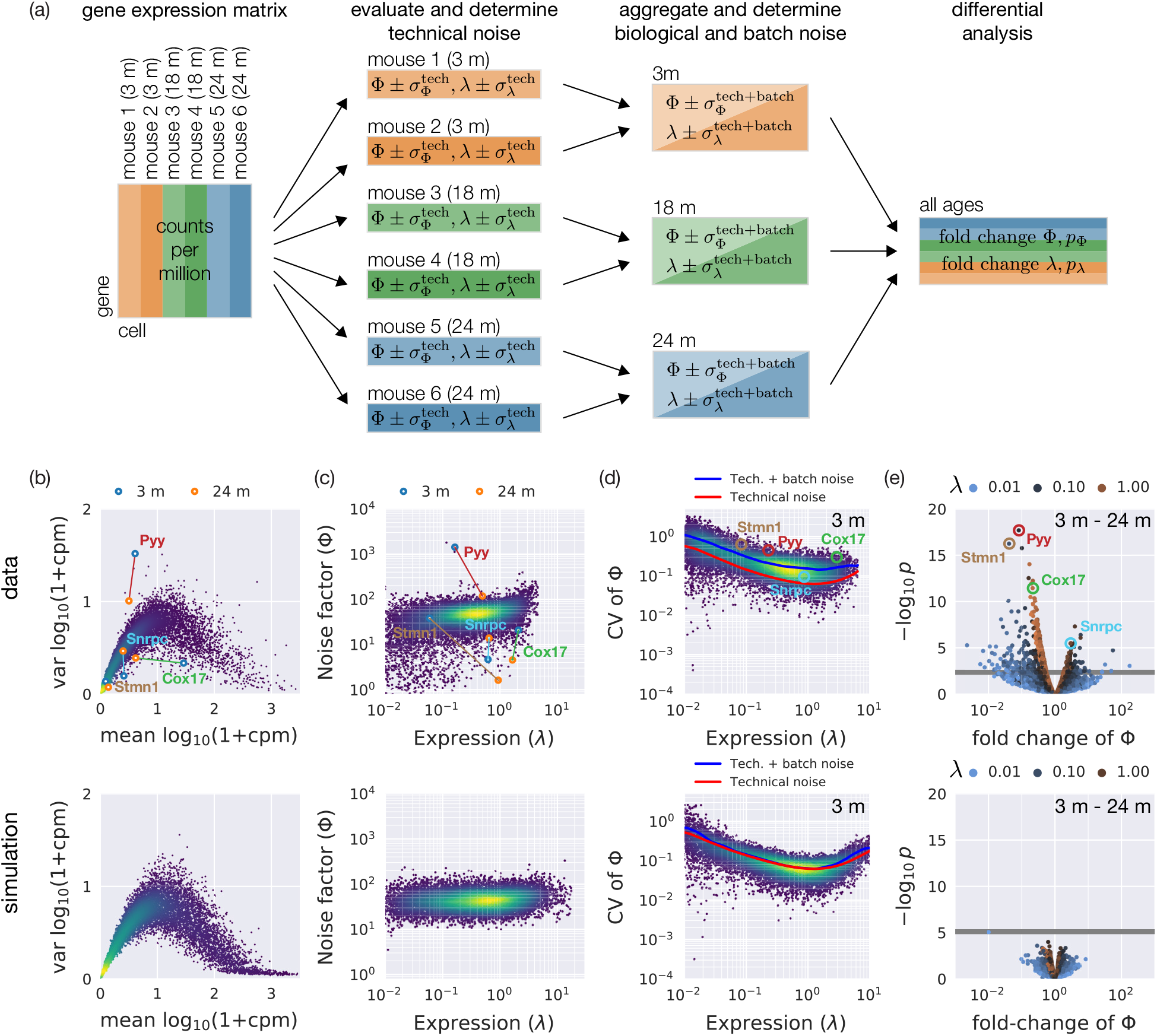
Pipeline to evaluate, aggregate, and perform differential analysis on expression and noise changes in aging mice. (a) The normalized gene expression matrix of one cell type, representing measured counts per million of each gene in each cell, is split by each mouse. The statistical model fits the expression *λ* and noise factor Φ to each gene, as well as their estimated uncertainties 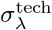 and 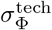. The model also determines a cell size variability of var *c* for each mouse. Results for different mice are aggregated by age. As each mouse has a different estimation of *λ* and Φ for each gene, mouse-to-mouse variation leads to an increased uncertainty 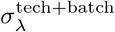 and 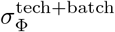, referring to potential batch effects. The total uncertainties are used in determining the adjusted *p* values of the fold change between ages. Genes with statistically significant changers in Φ are termed DNGs (differential noise genes). (b) Mean and variance of log_10_(1 + *x*) for each gene follow a characteristic shape that can be deconvolved into estimates of (c) expression (*λ*) and noise factor (Φ). (d) The measured coefficient of variation of Φ across cells isolated from different mice of the same age (3 m) is typically 2-3 times greater than the technical model estimates. We use the measured total uncertainty 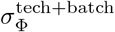 to test whether (e) fold changes between ages are statistically significant, corrected by Benjamini-Hochberg with *α* = 0.1. (Second row) All steps are validated by simulating gene expression with the statistical model and verifying that *p*-values are well calibrated and unbiased.

**FIG. S2:**
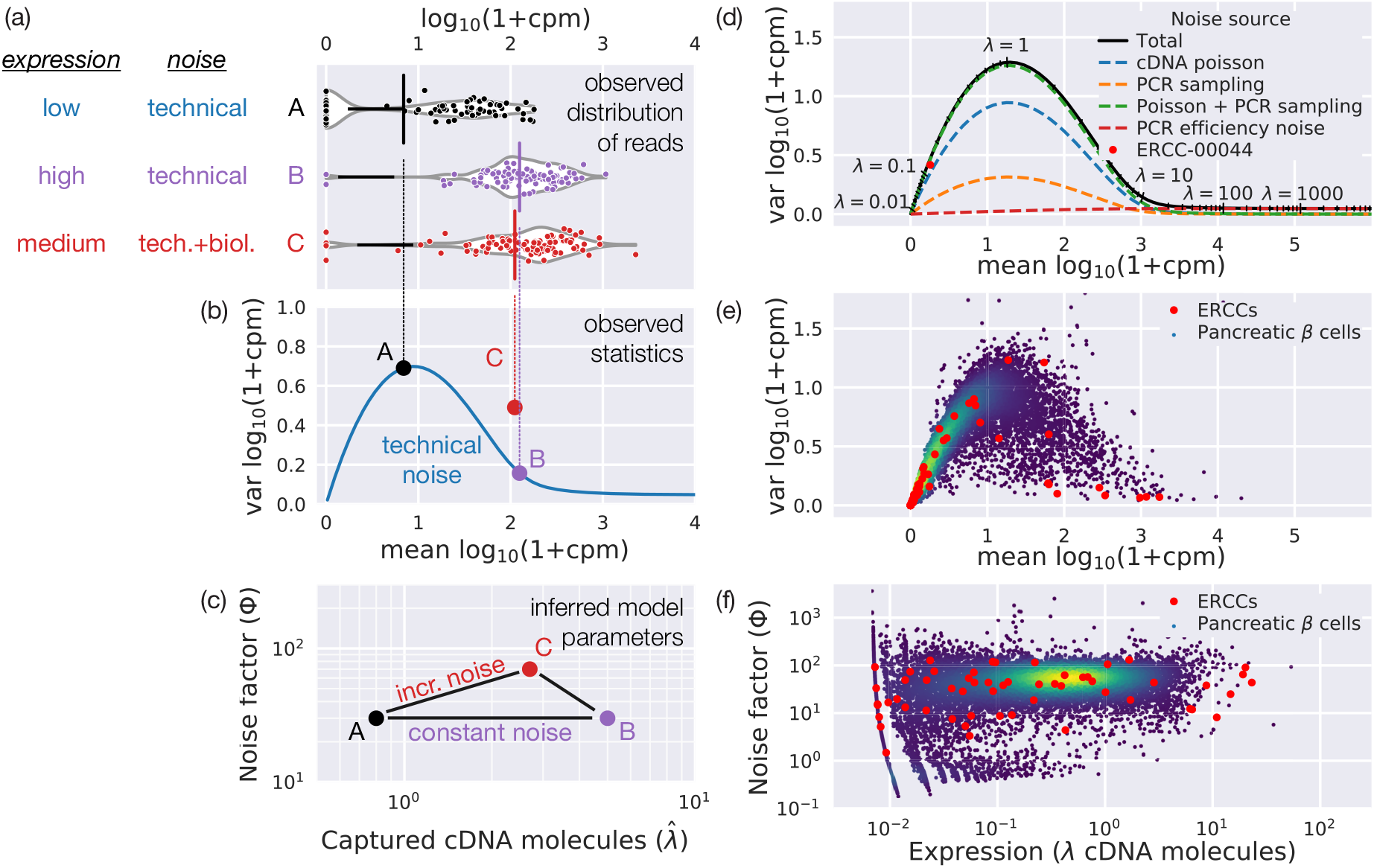
Changes in expression confound estimates of gene expression noise. (a) We simulate gene expression for one gene under three conditions: a change in expression from low (**A**, black) to high (**B**, purple) without biological noise, and moderate expression with biological noise (**C**, red). (b) With only technical (measurement) noise, changes in expression are accompanied by a substantial change in the measured variance due to technical noise alone (blue line). Biological gene expression noise increases the variance. (c) We fit an expression and noise factor to each condition to deconvolve changes in noise factor from changes in expression. (d) The noise model quantitatively predicts that the variance first increases and then decreases with expression (*λ*, black ticks). The counting statistics of cDNA capture dominated for *λ <* 10, while PCR efficiency noise dominates for *λ >* 20. (e) Mean versus variance of pancreatic *β* cell expression can be deconvolved into (f) *λ* and Φ that show little correlation over several orders of magnitude in expression. The majority of low expressed genes follow the same distribution as exogenous spike-in RNA (ERCCs, red points), which are expected to only have technical noise. The noise factor for most genes depends on their relative PCR efficiency.

**FIG. S3:**
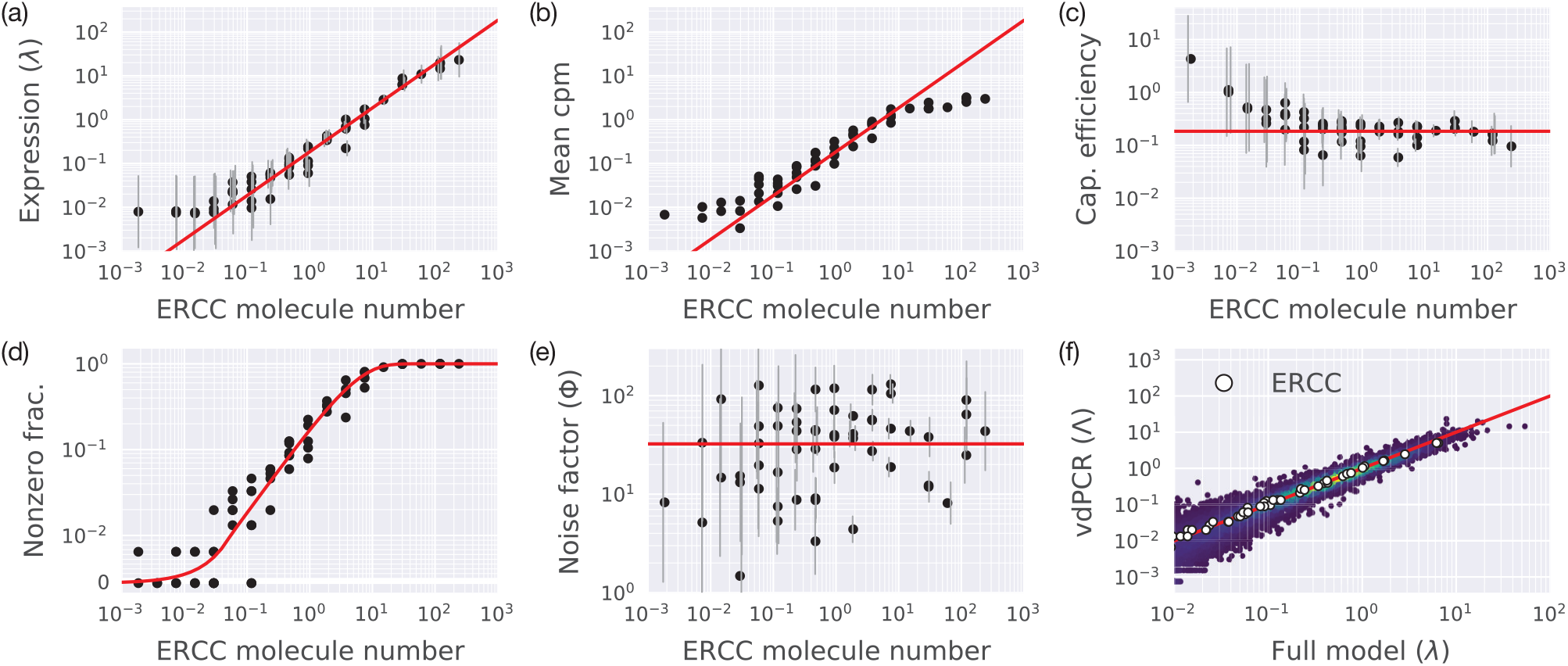
Validate model with spike-in RNA. Spike-in RNA (ERCCs) are used to validate the statistical model in pancreatic *β* cells. The ERCC molecule number is the expected number of molecules per well based on the known concentration and dilution. (a) Measured expression *λ*, the estimated number of cDNA molecules, is well correlated to the ERCC molecule number with a capture efficiency of 18.5(1.2)% (95% confidence), estimated from the ratio of expression to expected molecule number (curves in a-e and f are derived from this estimate). Vertical grey lines denoting 95% credible intervals are offset horizontally for clarity. (b) Mean counts per million, the conventional metric for expression in single-cell RNAseq, show a systematic decrease at high expression. (c) The estimated capture efficiencies for each control, *λ/mol*, is roughly constant for *>* 3 *×* 10^−2^ molecules. (d) Fraction of cells with nonzero expression of each control. The fit assumes a Poisson distribution with the measured capture efficiency. (e) Noise factor is independent of expression. The horizontal red line is the geometric mean of 32 cpm/cDNA molecule. In this context, the noise factor is equal to the PCR gain. Vertical grey lines denoting 95% credible intervals are offset horizontally for clarity and often significantly deviate from the mean, indicating that the fits can distinguish Φ for each control. (f) Full model fit of expression *λ* is strongly correlated with the vdPCR fit based only on dropout rates. White circles are ERCCs, red line is equality.

**FIG. S4:**
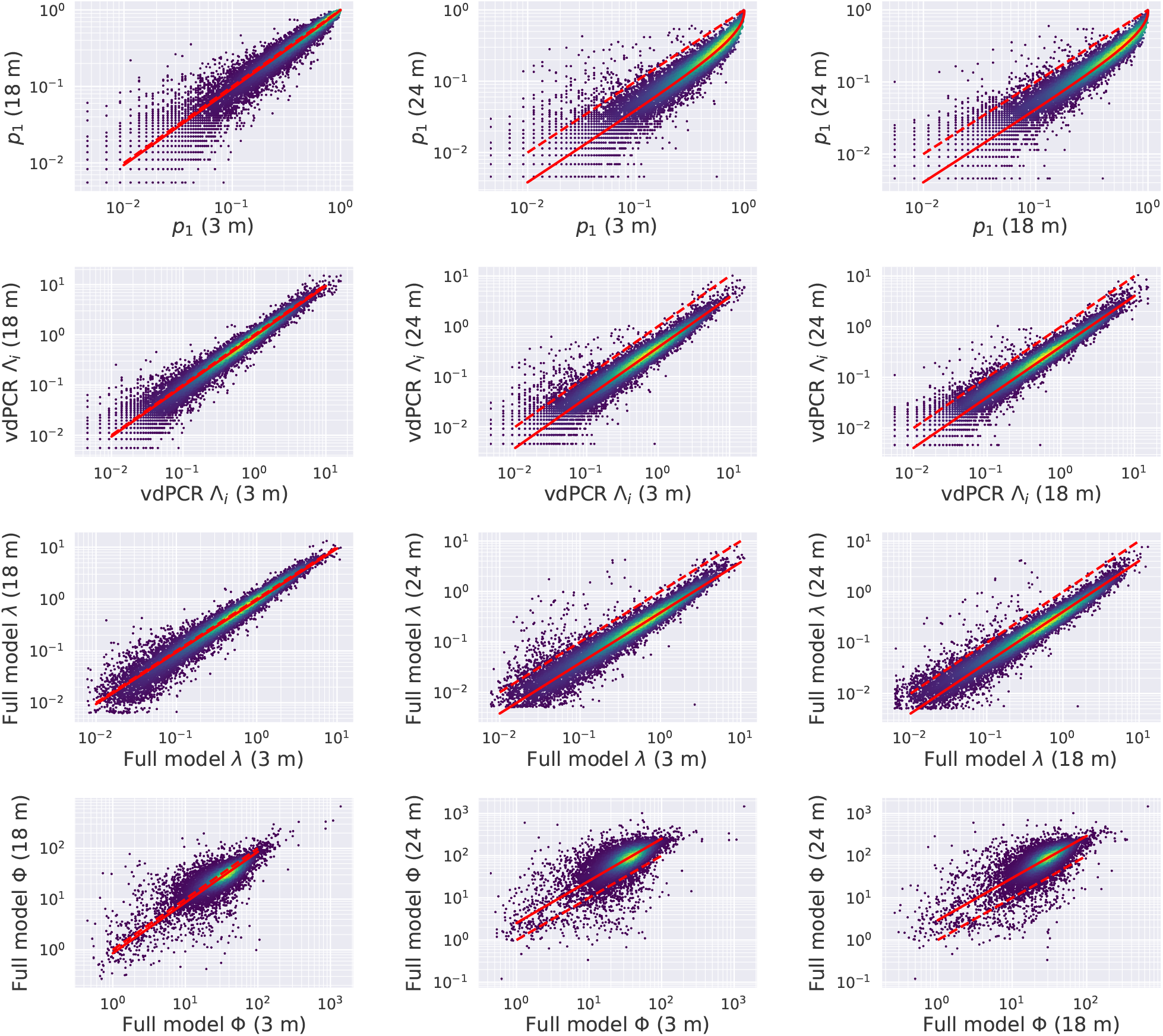
Number of cDNA molecules systematically decreases with age. Gene expression in pancreatic *β* cells isolated from 24 m mice shows a systematic decrease in the number of cDNA molecules per cell compared to cells isolated from 3 m and 18 m mice. This can be seen in the fraction of cells with nonzero count *p*_1_ (first row), the estimated expression from dropouts in vdPCR (second row), an estimated of expression from the full model (third row), and the noise factor (fourth row). The noise factor shows a systematic increase in 24 m mice because it is normalized by the total molecule number. Each point is one cell. Solid red curves are fits to the data, dashed curves show unity.

**FIG. S5:**
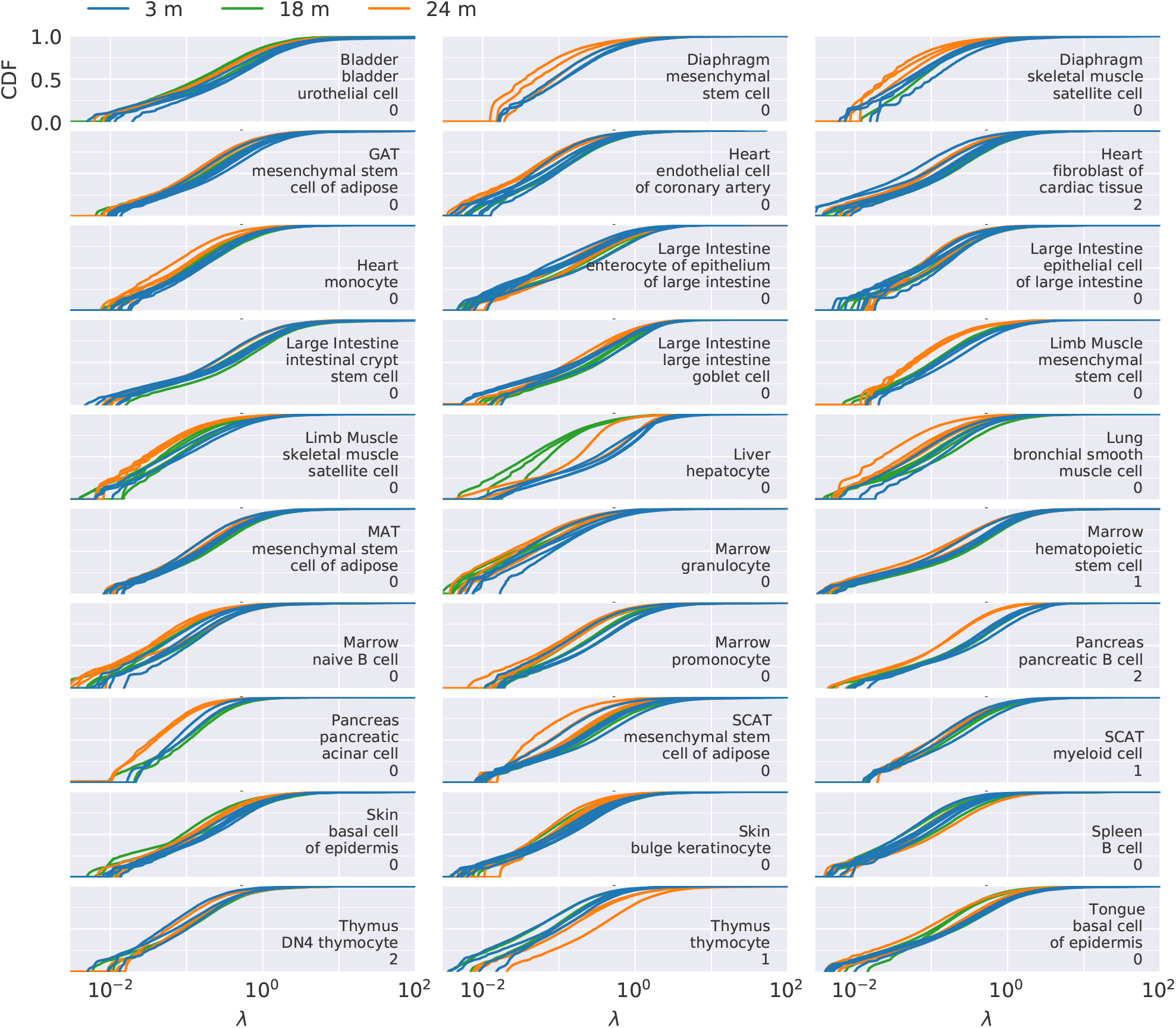
Distribution of expressions. Cumulative probability distribution of *λ*, the estimated number of cDNA molecules per cell, for genes with at least one read. Typically 80-97% of genes have low expression with less than one cDNA molecule (*λ <* 1). Exceptions include bladder urothelial cells (79%), intestinal crypt stem cells (77%), and pancreatic acinar cell (98%). Each line is the CDF of one mouse, colors represent age.

**FIG. S6:**
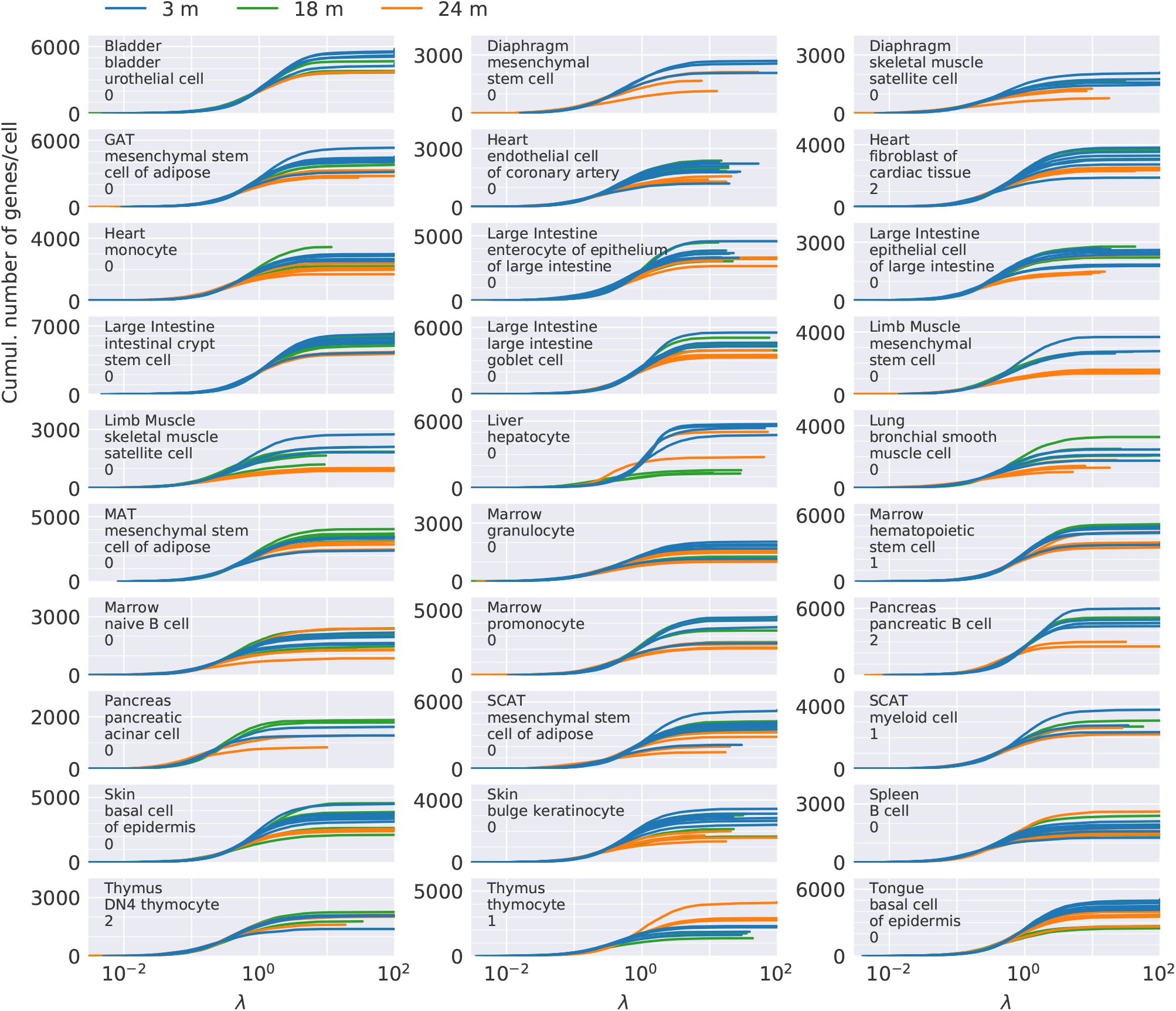
Distribution of unique number of genes per cell. Cumulative number of unique genes per cell (NOG) with expression less than *λ*, the estimated number of cDNA molecules per cell. While 80-97% of genes correspond to an expression of, *λ <* 1, these genes only make up of roughly half the number unique of genes seen per cell. Each line is the CDF of one mouse, colors represent age.

**FIG. S7:**
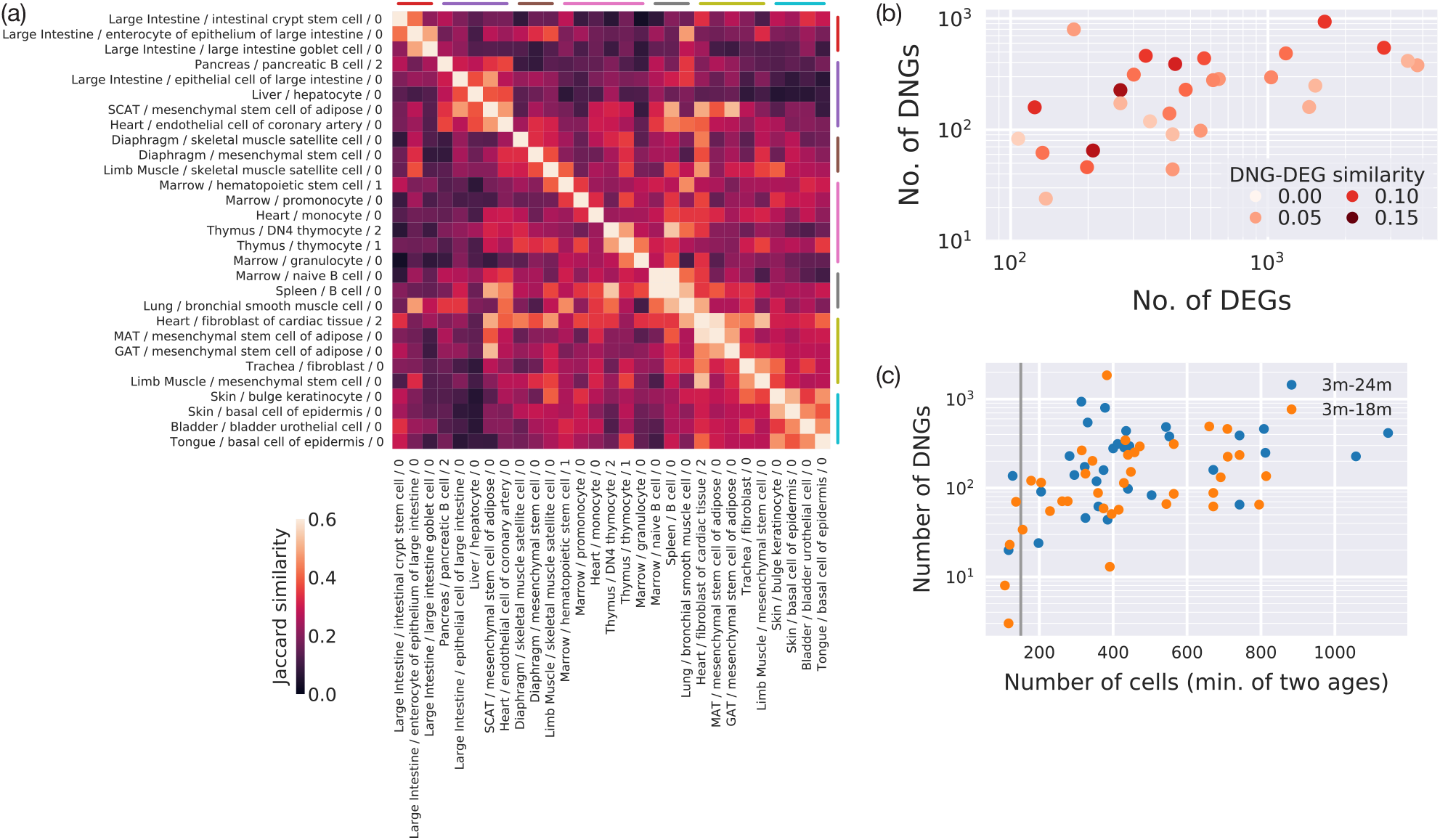
Cell types cluster based on similar DNGs. (a) Cell type are clustered by Jaccard similarity on the overlapping identity of DNGs. Some cell types have over 50% similarity. (b) The number of DNGs and differential expression genes (DEGs) is only weakly correlated. DNGs and DEGs typically have less than 10% Jaccard similarity within one cell type. (c) The number of DNGs discovered saturates slowly with the number of cells. A cutoff of 150 cells in each age removes cell types with insufficient statistics to find DNGs (vertical gray line).

**FIG. S8:**
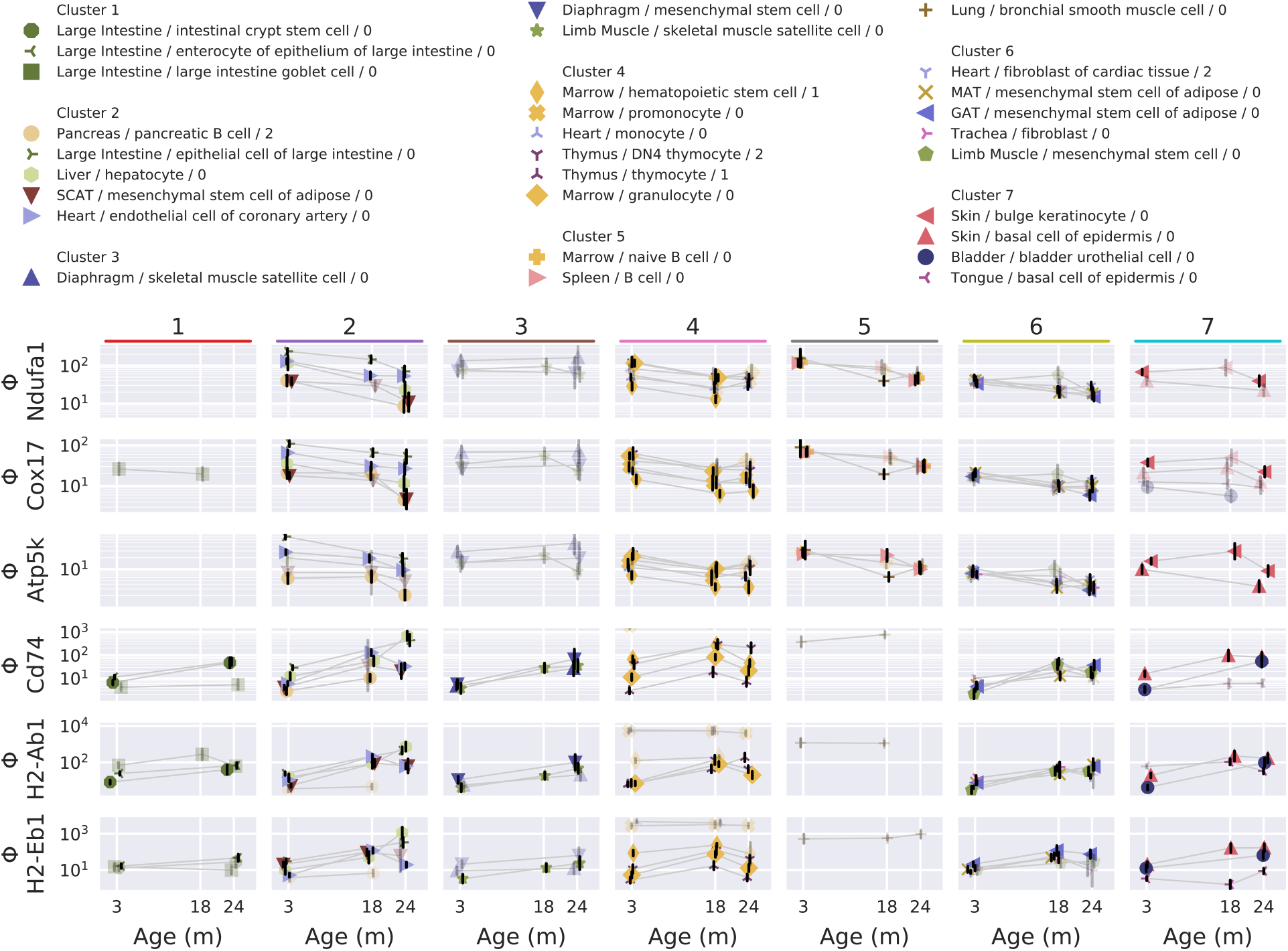
Change of noise factor with age for several differential noise genes. Select DNGs involved in oxidative phosphorylation (*Ndufa1, Cox17, Atp5k*) and antigen processing and presentation (*Cd74, H2-Ab1, H2-Eb1*) are grouped by cluster (Fig. S7).

**FIG. S9:**
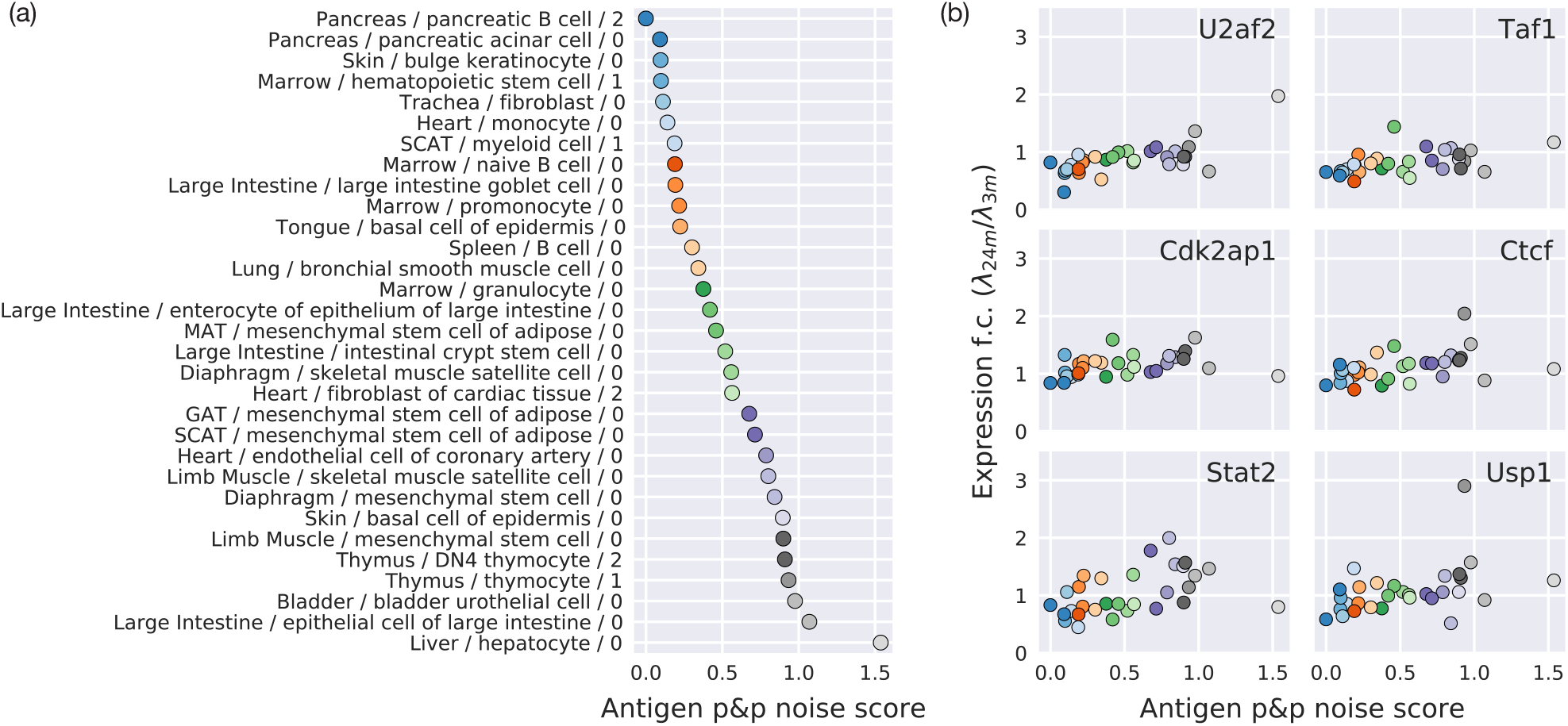
Increase in antigen processing and presentation is correlated with transcription factors and regulators. A noise score for antigen processing and presentation is correlated with increased noise in cells isolated from older mice. Similar to Fig. 5, top hits are predominately transcription factor (*Taf1, Ctcf, Stat2*), splicing factors (*U2af2*), cell cycle regulators (*Cdk2ap1*), and DNA damage repair regulators (*Usp1*).

## Supplemental Material

### 1. STATISTICAL MODEL

Our statistical of amplification and noise during single-cell mRNA sequencing has components that vary across cell (or more accurately well or droplet) *i* and genes *j*. For a pure population of cells, with noise limited only by technical noise involved in the capture and PCR, we expect the number of cDNA molecules in each well to arise from a Poisson distribution with mean *λ*_*j*_. Even if a cell expressed a fixed number of mRNA molecules in every cell, the relatively low probability of capture and reverse transcription would produce an approximately Poisson distribution.

Different cells have more or fewer total molecules of cDNA due to differences in capture efficiency, cell size, degradation, lysis efficiency, etc. by a relative amount *c*_*i*_, where the mean is defined to be 1 and the variance is measured to be var *c*_*i*_ = 0.1 − 0.3. Some cell types, such as hepatocytes, show extreme examples of up to 2.0. To account for this effect, we model the number of cDNA molecules of gene *j* on well *i* as *n*_*ij*_ *∼* Poisson(*c*_*i*_*λ*_*j*_), with a Fano factor *F* = 1 + *λ*_*j*_var *c*_*i*_ greater than 1, but only slightly so for low expressed genes (*λ*_*j*_ ≪ 1). The parameter *c*_*i*_ is equivalent to *β* in Ref. S13, however, we find that virtual digital PCR is a superior method to estimate *c*_*i*_ from low-expressed genes, and use spike-in controls to validate the estimate.

Ideally, we would measure the distributions *n*_*ij*_ with single molecule counting, but this is difficult to perform for many genes and cells simultaneously. Instead, we use data processed through the SmartSeq2 pipeline generated by the Tabula Muris Senis collaboration. On each well, cDNA is amplified through typically 12 stages of PCR, with downstream steps of library preparation, sequencing, alignment, and other computational analysis to generate a counts table. The stochastic nature of PCR, by which a molecule has a finite probability of being replicated in each cycle, adds a noise term *y*_*ij*_ dominated by the first few cycles of replication and derived below in Sec. I B. In brief, we approximate *y*_*ij*_ as a truncated Gaussian with variance *n*_*ij*_*/*3, the worst-case scenario corresponding to a PCR efficiency of *p*_*j*_ *≈* 50%.

After many cycles, a single molecule of cDNA will be converted into 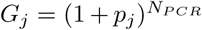 counts per million for gene *j*. The number *G*_*j*_ typically varies from 10 to 100 cpm in cells isolated from 3 m mice. Slight well-to-well variations causes differences in PCR efficiency from well to well, leading to *g*_*ij*_ that is expected to be log-normally distributed around *G*_*j*_. While the source of variation here is unknown, it’s expected that small, normally distributed variations in *p*_*j*_ would lead to a log-normal distribution in *g*_*ij*_. In addition, variations in total cDNA leads to ratio of *g*_*ij*_*/c*_*i*_ of normalized reads per molecule cDNA, so the measured normalized reads is *x*_*ij*_ = (*g*_*ij*_*/c*_*i*_)(*n*_*ij*_ + *y*_*ij*_). The terms are summarized below, where *𝒩* (*μ, σ*^2^) denotes the normal distribution with mean *μ* and variance *σ*^2^.

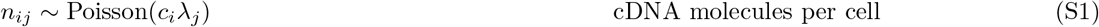

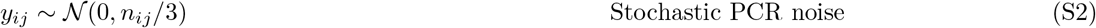

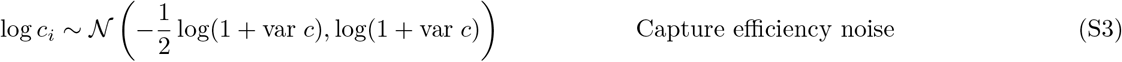

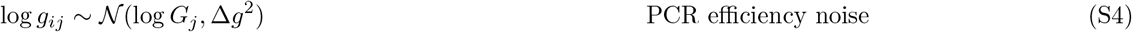

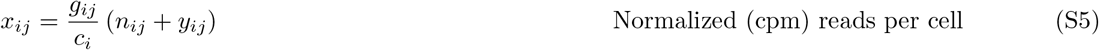

The amplification process increases the Fano factor. If capture efficiency noise is neglected, then the variance and mean of *n*_*ij*_ are both *λ*_*i*_. Capture efficiency noise increases the variance of *n*_*ij*_ to *λ*(1 + *λ* var *c*), which is negligible for small *λ*. Stochastic PCR increases the variance by 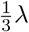. After amplification by a factor *G*, the mean and variance increase by *G* and *G*^2^, respectively, causing the final Fano factor (variance over mean) to increase by *G*. For purely technical noise, we expect a Fano factor of (4*/*3)*G*(1 + *λ* var *c*). For purely technical noise, with a nearly noiseless distribution of mRNA, we define a noise factor Φ = *G*.

Biological variation is expected to increase the Fano factor of cDNA. If the true Fano factor of cDNA molecules is *F*, due to biological variation, then the final Fano factor is approximately (4*/*3)*FG*_PCR_(1 + *λ* var *c*), where *G*_PCR_ is the counts per million per cDNA molecule after PCR, library preparation, sequencing, alignment, etc. The actual impact on the measured mean and variance depends on the biological distribution of RNA. In brief, we fit data to the above model to get estimates of 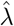 and 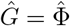, where Φ is the noise factor and includes both the PCR gain and a biological noise term.

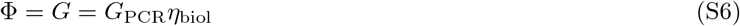

We assume that *G*_PCR_ is constant across different ages and conditions, so that changes in Φ arise only from changes in the true biological noise *η*_biol_. This parameterization has the advantage that it is model independent, as the precise form of *η*_biol_ depends in detail upon the specific distribution of the biological variation (see Sec. I G for further discussion).

#### A. Capture efficiency noise and virtual digital PCR

Differences in cell sizes, availability of mRNA, capture efficiency, or reverse transcription processes, will increase the observed cell to cell variability in cDNA. Unaccounted-for changes may be confused with biological gene expression noise, which similarly changes the observed cell-to-cell counts in a given gene. For this reason it is crucial to understand and calibrate cell size variability to distinguish it from more interesting biological effects. Fortunately, we can distinguish and calibrate cell size variability from the binarized count table. If the sequencing read depth is sufficiently large, as is the case of the *Tabula Muris Senis* experiment, then we expect we are very likely to see at least one read when there is at least one molecule of cDNA for a given transcript.

While the distribution of reads may be distorted by PCR, we can calculate a binarized count table that encapsulates capture noise and cell size variability. We define a binarized count table *z*_*ij*_ for gene *i* and cell *j* as,

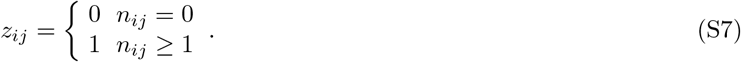

The likelihood of measuring a particular value of *z*_*ij*_ is,

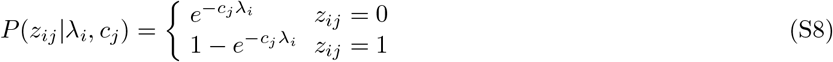

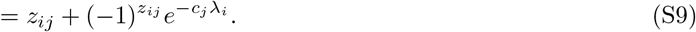

We fit the sets {*λ*_*i*_} and {*c*_*j*_} by minimizing the log likelihood function log ℒ = Σ_*i,j*_ log *P*(*z*_*ij*_|*λ*_*i*_, *c*_*j*_). The likelihood ℒ is invariant under the rescaling *λ*_*i*_ *→ αλ*_*i*_ and *c*_*j*_ *→ c*_*j*_*/α*, so we can additionally require ⟨*c*_*j*_ ⟩ = 1. The problem is considerably overdetermined with *n*_cells_ + *n*_genes_ − 1 fit parameters, far less than the *n*_cells_ *× n*_genes_ observations.

In practice, we first find the solutions 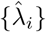 that minimize the log ℒ for fixed *c*_*j*_ = 1, then fix 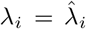 to solve for {*c*_*j*_}. We iterate between minimizing log ℒ for fixed {*c*_*j*_} or {*λ*_*i*_}, each time fixing the variables to the solution from the previous iteration 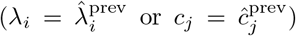. We continue until the maximum error is less than 10^−6^ 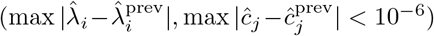. We observe that the error decreases by approximately an order of magnitude each iteration and reaches this threshold after six to eight iterations. After each iteration we normalize the fits to ensure ⟨*c*_*j*_⟩ = 1 (rescaling *λ*_*i*_ *→ λ*_*i*_ *c*_*j*_ and ⟨*c*_*j*_ ⟩→*c*_*j*_/ ⟨*c*_*j*_⟩). We did not observe better performance by analytically supplying the first and second derivatives of the log likelihood function or solving for the root of the first derivative. In future work, the minimization should be performed with priors reflecting the distribution of *λ*_*i*_ and *c*_*j*_, for instance with an empirical Bayes method.

Since *z*_*ij*_ can only be 0 or 1, its variance (averaged across all cells *j* for a particular gene *i*) is var_*j*_ *z*_*ij*_ = *p*_*i*_(1 *− p*_*i*_), where *p*_*i*_ is the fraction of nonzero *z*_*ij*_ for a given gene 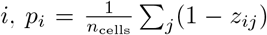 (Fig. 2c, blue points). We use the fits *ĉ*_*j*_ to estimate 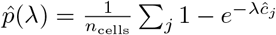 (red solid line is var 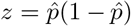. If we neglect capture efficiency noise, then 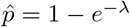 and var *z* = *e*^*−λ*^(1 *− e*^*−λ*^) (black dashed line).

capture efficiency noise increases the variance of the true cDNA molecule number *n*_*ij*_. Since *n*_*ij*_ ∼ Poisson(*λ*_*i*_*c*_*j*_) it can be shown that that the Fano factor is increased beyond the Poisson prediction of one.

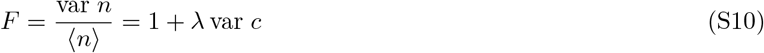

where var *c* is taken over all *c*_*j*_ and ⟨*c* ⟩ = 1.

#### B. Stochastic PCR noise

After each PCR cycle, there is a probability *p* of each molecule getting replicated. If there are *m* molecules before a cycle of PCR, the probability of ending up with *m* molecules is 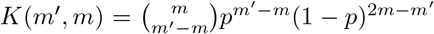. We can then calculate that a probability distribution function *pdf*_*n*_(*m*) after *n* cycles of PCR is transformed into *pdf*_*n*+1_(*m*) after the *n* + 1 cycles of PCR as,

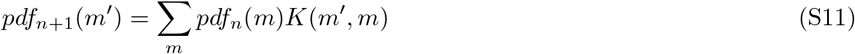

After many cycles, the numbers of molecules *m* because substantial and allows us to make a continuum approximation. If we start with *m* molecules, we expect that the resulting molecule number distribution has a mean of (1 + *p*)*m* and a variance of *mp*(1 *− p*), and so we make a gaussian approximation with the corresponding mean and variance.

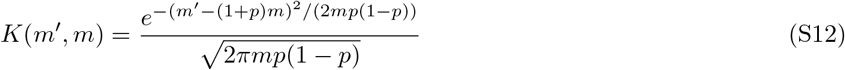

After *N* PCR cycles, the distribution’s mean increases by (1 + *p*)^*N*^. Since our goal is to infer the input number of molecules, it is most helpful to work with an input-referred distribution in terms of the variable *x* = *m/*(1 + *p*)^*N*^. In other words, we normalize the distribution after each cycle. Now the mean of the distribution about *x* does not change after each PCR cycle, and we will see the variance saturates. Substituting *m* = *x*(1 + *p*)^*N*^ and *m* = *x*(1 + *p*)^*N*+1^,

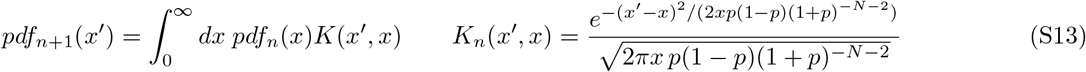

After each cycle, this convolution increases the variance by *μ*_*n*_*p*(1 *− p*)(1 + *p*)^*−N*^. If the initial variance of *pdf*_0_(*x*) is 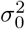, then after *N* cycles the variance is,

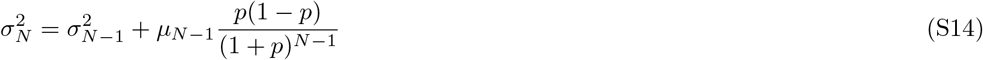

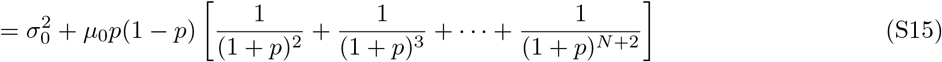

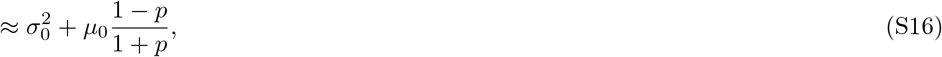

where we used the fact that *μ*_*n*_ = *μ*_0_ is constant and assumed *n* is large. The first term 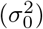 is the input noise, and the second term is the additional technical noise added by the PCR amplification process. If the initial input is a Poisson distribution with 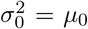, then the variance of the Poisson distribution exceeds the stochastic noise of the amplifier, since (1 *− p*)*/*(1 + *p*) *<* 1. When *p* = 1*/*2, the input-referred noise is *μ*_0_*/*3, and the final (input plus added noise) variance is 4*μ*_0_*/*3. The exact shape of the distribution can be calculated with the series,

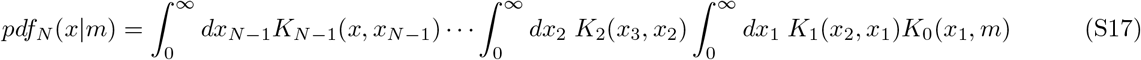

Integrating this series is nontrivial. When *m* ≫ 1, we can make a gaussian approximation as before,

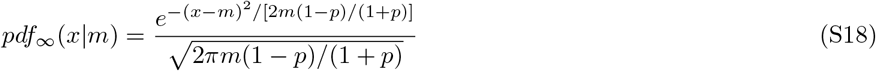

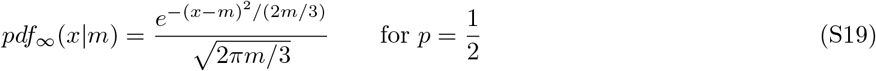

#### C. Comparison to previous work

Previous work has elucidated the role of capture efficiency noise by counting cDNA molecules. The most common approaches are cDNA barcodes, typically termed unique molecular identifiers (UMIs), and single-molecule fluorescence *in situ* hybridization (smFISH). UMIs remove PCR distortion and bias by identifying which reads were amplified from the same, original molecule. For UMIs to provide faithful counting there must be sufficient reads per UMI to minimize dropout, including for transcripts which suffer from low PCR gain (low PCR bias). The primary tradeoff of UMIs is that the additional tagging step lowers the likelihood of reverse transcription and thereby decreases the efficiency of molecule counting. smFISH techniques count molecules directly with fluorescent probes, foregoing PCR amplification.

**FIG. S10:**
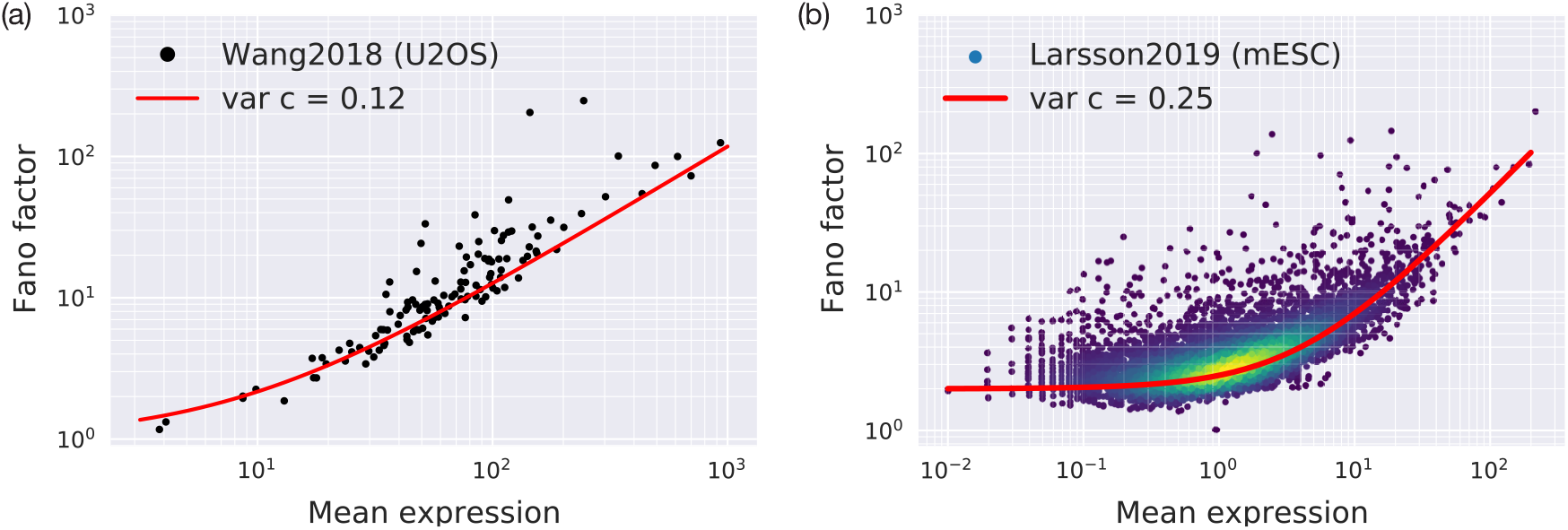
Fano factor versus mean expression in previous work. Mean expression is the estimated number of molecules, where molecule counting is estimated from (a) merFISH and (b) unique molecular identifiers added before amplification and sequencing. In both cases, the majority of genes follow the expected theory of a Poisson distribution plus a capture efficiency or cell size variability, Eq. S10.

Several previous papers have quantitatively studied the relationship of variance and mean of cDNAs [S13, S24, S38]. Ref. S24 used expansion microscopy and merFISH to count highly expressed transcripts in a relatively uniform cell population (U2OS). We observe excellent agreement with Eq. S10 in this orthogonal technique (Fig. S10a).

Ref. [S38] has several unusual features worth considering. This paper focuses on single-cell expression of heterozygous alleles in crossbred mouse embryonic stem cells. The resulting distributions are fit to a standard two-state model of bursting gene expression [S28, S30]. Surprisingly, the paper neglects technical noise, despite previous work demonstrating that the majority of the noise is technical [S14]. Bursting gene expression is typically characterized by four rate constants, *k*_on_ and *k*_off_ rates switching between transcriptionally active and inactive state, a rate of transcription *k*_syn_ in the active state, and a transcript degradation rate *δ*. Single single-cell RNAseq captures snapshots of gene expression in time, so only dimensionless parameters are measurable: *α* = *k*_on_*/d, β* = *k*_off_/*δ*, and *s* = *k*_syn_*/d*. The distribution of a gene is given by *p∼* Beta(*α, β*) and *n ∼* Poisson(*sα*) [S30], with a mean *μ* and Fano factor *F* of this model are as follows.

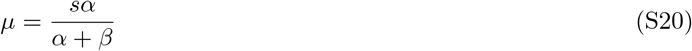

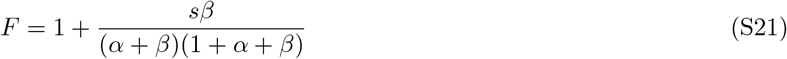

The distributions is similar to technical noise (Eq. S1 and S4) and produces distributions that are qualitatively indistinguishable from it when *α, β* ≪ 1. Only careful calibration of the parameters, particularly var *c*, can quantitatively distinguish the two. The challenge of discerning biological bursting gene expression from technical noise in single-cell gene expression has long been recognized [S14, S29, S30].

Estimating the Fano factor from *α, β*, and *s* (Eq. S21) allows us to test the data against of model of capture and capture efficiency noise (Eqs. S1, S3, and S10). We observe excellent agreement with *F* = 2(1 + *λ*var *c*), where var *c* = 0.25, strongly suggesting that the majority of the dataset is well explained by technical noise alone [S30]. The capture efficiency noise parameter var *c* = 0.25 is consistent with that observed in *Tabula Muris Senis* and slightly larger than for merFISH. Surprisingly, the Fano factor is approximately two for even low expressed genes. This can be most parsimoniously explained by each transcript being counted, on average, twice.

#### D. Fitting the model

The parameters of the model are *λ*_*j*_ and *G*_*j*_ for each gene and var *c* and Δ*g* for all genes. We fit this model to each gene, in each mouse, for each cell type. While many methods exist to estimate *λ*_*j*_ and *G*_*j*_, such as maximum likelihood estimation and Bayesian estimation, we simply fit summary statistics, which allows for a substantial increase in computational speed at an acceptable sacrifice in precision. We measure the mean *m*_*j*_ and variance *v*_*j*_ of log_10_(1+*x*_*ij*_), averaged over the *i* cells within one mouse and cell type, and solve for the theoretical *m*(*λ*_*j*_, *G*_*j*_, var *c*, Δ*g*) and *v*(*λ*_*j*_, *G*_*j*_, var *c*, Δ*g*) such that 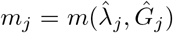 and 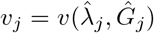 for each gene *j*, matching the measured mean and variance against their theoretical estimates. This is done for fixed var *c* and Δ*g* for all genes. The functions *m* and *v* are calculated from a numerical integration of Eqs. S1-S4. We will drop the indices *i* and *j* for convenience.

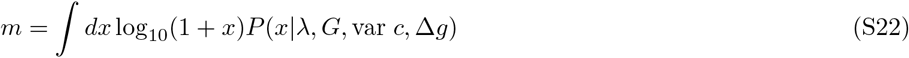

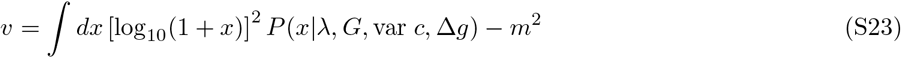

The probability distribution function *P*(*x*|*λ, G*, var *c*, Δ*g*), as derived from Eqs. S1-S4, involves integrating over all *g, y, c*, and summing over *n*. The special case of *x* = 0 is handled separately.

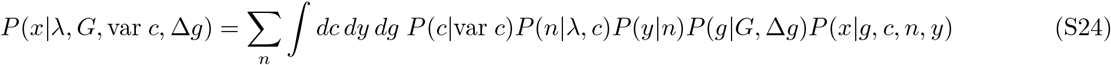

The computational challenge is to invert the quadruple integral/sum in Eq. S24 and Eqs. S22-S23 to determine 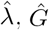, *Ĝ* for each of approximately 10,000 genes in the approximately 100 cell types across about a dozen mice.

#### E. Numerical approximations of the PDF

Calculating Eq. S24 across many genes is made tractable through caching intermediate results and careful numerical sampling. First, we rewrite the calculation in terms of *u* = log(*x/G*), which removes *G* from the calculation. We add in only at last step *m*(*λ, G*) = Σ_*n*_ ∫ *dy dg dc* log_10_(1 + *Ge*^*u*^)*P*(*u*|*c*, var *c*, Δ*g*), and similarly for *v*(*λ, G*). For a fixed Δ*g* and var *c* for each dataset, we cache results for *P*(*u*|*n*), given an initial number of molecules *n*, following Eqs. S5,S2, and S4.

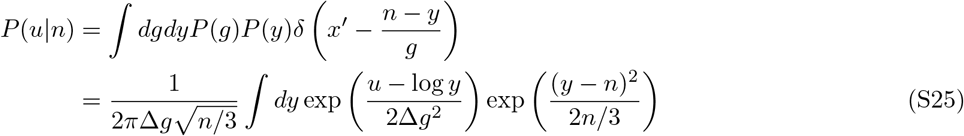

This equation can be solved efficiently by Gaussian-Hermite quadrature, by which an integral may be approximated as 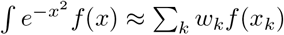, where *x*_k_ is the *k*^th^ root of the Hermite polynomial and *w*_*k*_ is the weight. Here, we use 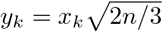.

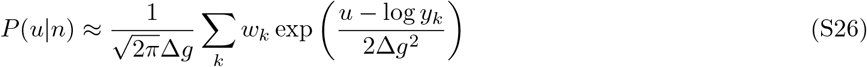

We find an excellent tradeoff between computational efficiency and accuracy with only 20 roots, and sampling *u* from − 4 ln 10 to 8 ln 10 in 1,000 steps, which samples *x*′ from 10^−4^ to 10^8^. Finding the appropriate grid to cache *n* is more subtle. If Δ*g* = 0, the integral becomes trivial and *n* is sampled densely with linearly increasing spacing that ensures that a Poisson distribution is samples with a constant number of points for all *λ*.

Once *P*(*u*|*n*) is cached, its values can be looked up to find *P*(*u*|*λ*, var *c* = 0) from Eq. S1. In principle, one would want to calculate

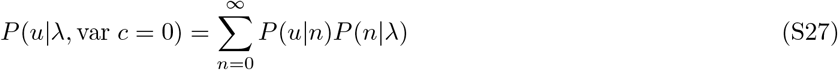

However, *n* is sampled along a grid for efficiency, and so many integer numbers are skipped. Instead, we calculate,

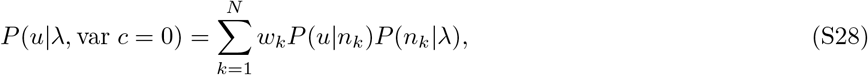

where 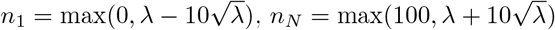, and *w*_*k*_ = 1*/* Σ_*k*_ *P*(*u*|*n*_*k*_).

Finally, capture efficiency noise in Eq. S3 is computed. Since we assume that *c* is log-normally distributed, we calculate Gaussian-Hermite quadratue of the normally distributed *v* = log *c*. Since the mean of *c* is defined to be one, then the variance of *v* is *σ*^2^ = log(1 + var *c*) and mean is 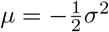. Finally, each *x* is normalized by that cell’s total reads, *x*′ = (*n* + *y*)*/c*, so we need to calculate sum across *P*(*u*′ − log *c*|*cλ*).

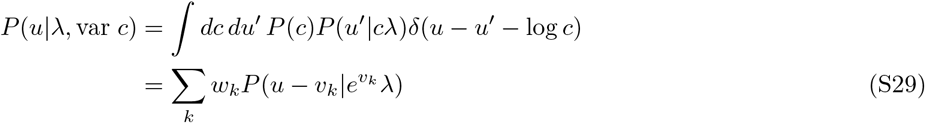

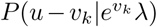 is directly calculated from Eq. S28, and *v*_*k*_, *w*_*k*_ are derived from the roots of the Hermite polynomial and weights of Gauss-Hermite quadrature, respectively. We find 10 roots are sufficient for good precision. Eq. S24 is found by substituting *x* = *Ge*^*u*^. The expected mean *m*(*λ, G*, var *c*, Δ*g*) and variance *v*(*λ, G*, var *c*, Δ*g*) can be computed directly. For typical parameters, calculating Eq. S29 requires 250 ms each iteration, summing over *n* (Eq. S28) and *v* (Eq. S29) from precomputed values of Eq. S26. Further discussion of uncertainties in estimating *λ* and *G* from a given distribution are discussed in Sec. I H.

We note that, without normalization, Eq. S29 predicts a very substantial increase in variance from capture efficiency noise. Normalizing by counts per million, *x*_*ij*_ = (*g*_*ij*_*/c*_*i*_)(*n*_*ij*_ + *y*_*ij*_), substantially suppresses this effect, validating our use of counts per million normalization. Other normalization schemes may not directly benefit from this effect.

#### F. Changes in expression confound technical noise estimates

One of the primary challenges of estimating changes of biological gene expression variation between two samples is that the technical noise depends on the mean gene expression level, which often varies substantially with age and experimental conditions. A simulation of the noise model in Fig. S2 highlights the effect. Since PCR produces long-tailed distributions, expression is typically transformed as “log plus one” to compress long-tailed outliers while retaining dropouts, cells with zero expression. We consider three possible distributions of the same gene (Fig. S2a). As the gene expression *λ* increases from low (A, black points) to high (B, purple points), the measured variance decreases (Fig. S2b, an effect caused by technical noise and by the log plus one transformation). When biological variation broadens the distribution of cDNA in a cell, we expect to measure a distribution with increased variance such as C (red point), which has the same variance as A but a greater variance than expected for technical noise at this expression (B and blue line). Our model allows us to transform the data to the estimates 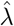, the mean number of cDNA molecules per cell, and 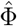, the relative noise factor (Fig. S2c). The transformed gene expression allows us to compare changes in noise and expression independently.

#### G. Impact of biological noise

The above analysis is used to fit genes limited by only technical noise to the probability distribution described in Eq. S24. We then heuristically define an estimated noise factor 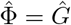 that captures both the PCR amplification and potential biological variation in the data. This heuristic is chosen because biological variation increases the variance of reads indistinguishable from an increase in PCR amplification, as they both increase the final, measured, Fano factor. Simulations based on several biological noise models are shown in Fig. S11, demonstrating that a negative binomial distribution and Poisson mixture distribution both prod. uce data that can be fit by an effectively greater *NF*. We distinguish biological variation by searching for (1) differential variance genes, where *NF* changes significantly between two groups, such as different ages, or (2) high variance genes, where *NF* greatly exceeds the expected range of *G* calibrated by low expression, low noise genes.

**FIG. S11:**
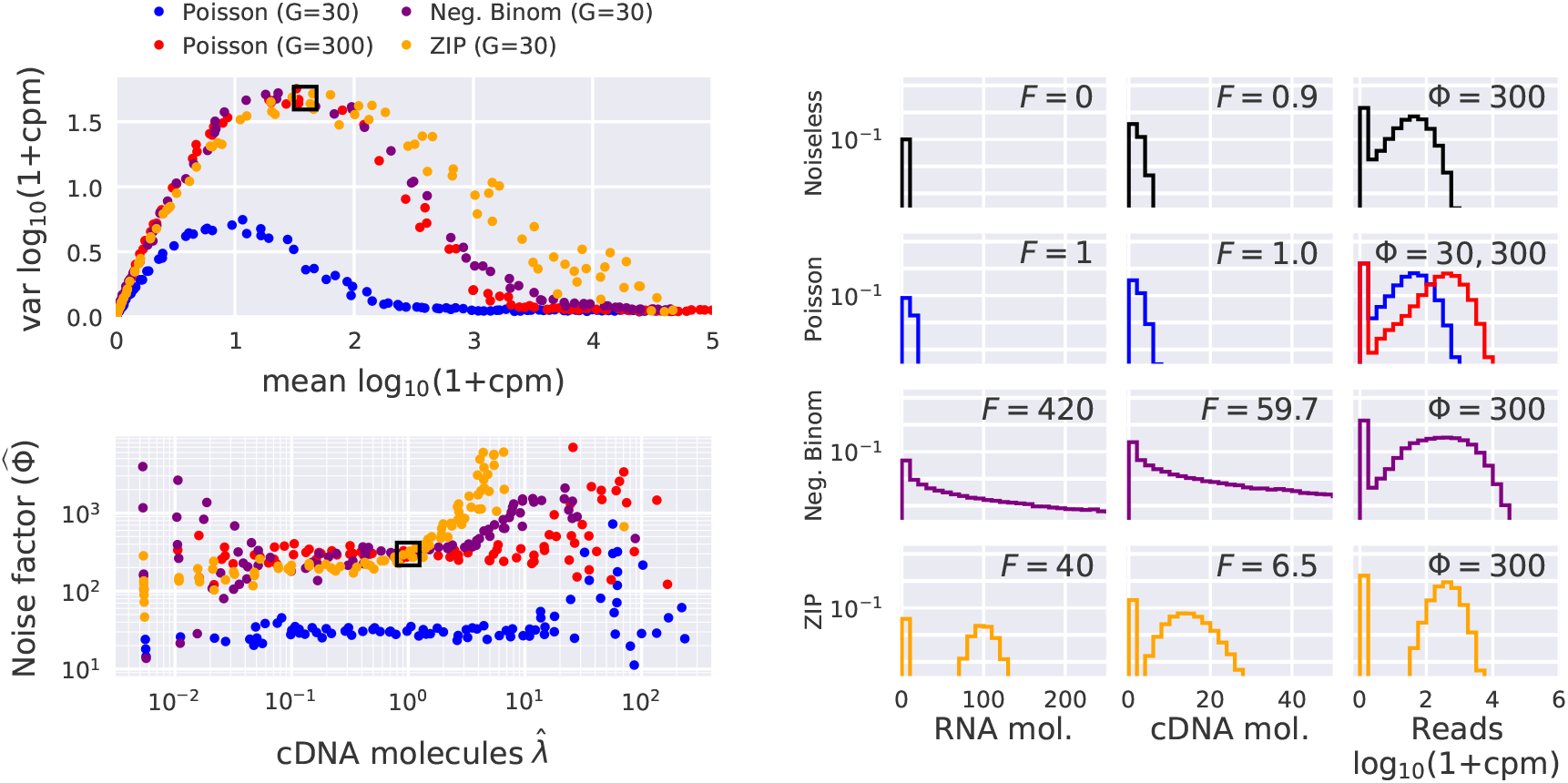
Simulation of noise factor extraction for several forms of biological noise. We simulate the negative binomial distribution (purple) and zero-inflated Poisson (ZIP, orange), which are two limits of bursting gene expression [S28–S30]. (a) The variance vs mean relation is similar to a Poisson distribution with increased *G* (blue and red). (b) Estimating the noise factor Φ and the expression *λ* generally deconvolves changes in expression with that of noise, though the accuracy depends on the specific biological model. Φ correctly measures that the biologically noise distributions have increased noise as compared to a Poisson distribution with the same PCR gain (c) Theoretical distributions of the RNA molecule number, cDNA molecule number, and reads for each distribution.

The technical noise model (Eqs. S1-S4) can be extended to capture these effects. The distribution of cDNA molecules depends on unknown parameters *θ*_1_, …, *θ*_*k*_ that, for a pure population, do not depend on the cell *j*. Eqs. S1 can be replaced by the following term.

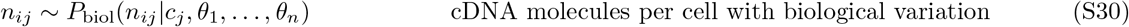

The measured distribution of reads, normalized by counts per million, now depends on

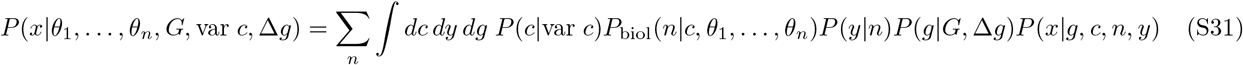

In the technical noise model, we assumed that the original distribution of RNA molecules is fixed, which implies that the subsampled distribution of cDNA molecules is a binomial distribution, which can be well approximated by a Poisson distribution with *λ*^cDNA^ = *pλ*^mRNA^, where *p* is the probability that one molecule of mRNA successfully is reverse transcribed and amplified. The Poisson approximation allows us to model the cDNA distribution without knowledge of *p*. capture efficiency noise (total mRNA content variation) makes the sampling probability dependent on the specific cell, with a mean number of cDNA molecules *cλ*^cDNA^ = *cpλ*^mRNA^ for a relative capture efficiency *c*. For an arbitrary probability distribution of mRNA molecules ℱ(*m*; *θ*_1_, …, *θ*_*k*_), subsampling the distribution of mRNA may be more complex. We note that if *F*_mRNA_ is the Fano factor of the mRNA molecule distribution, then we note that *F*_cDNA_ = 1 + *p*(*F*_mRNA_ − 1) and *NF ≈ GF*_cDNA_ [1 + *p*(*F*_mRNA_ − 1)]. We can then calculate the distribution of *n* cDNA molecules from *m* is the number of mRNA molecules in a cell with relative size *c*.

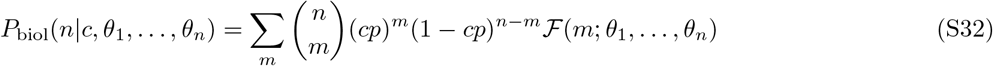

As before, Eq. S31 can be integrated to find the mean *m, v* and fit to the observed mean and variance of each gene.

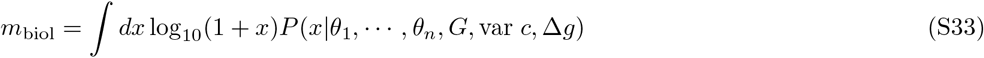

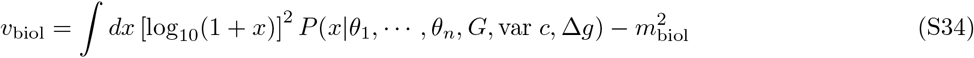

In principle we can fit these new parameters to the technical noise model with *m*_biol_ = *m*(*λ*^*∗*^, *NF* ^*∗*^) and *v*_biol_ = *v*(*λ*^*∗*^, *NF* ^*∗*^) for effective values of *λ*^*∗*^ and *NF* ^*∗*^, where we expected that *NF* ^*∗*^≳ *G*, with *NF* = *G* when there is no biological variation. Example of different *F* are shown in Fig. S11.

#### H. Probability of measuring a particular mean and variance

Once a *P*(*x*|*λ, G*) from Eq. S24 is numerically calculated, we want to know the uncertainties of an observation. Here we will use a frequentist approach for computational convenience. The conditional probability *P*(*m*′, *v* ′|*λ, G*) of measuring mean *m* and variance *v* of log_10_(1 + *x*), given a true *λ, G*, depends on the vector 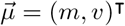 of the mean values, and the vector 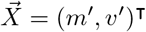 of possible values,

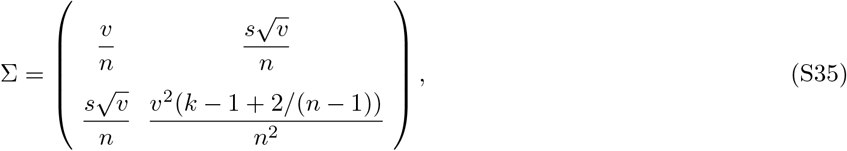

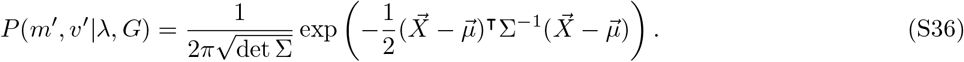

We would also like to determine the uncertainty in determining *λ, G* for each gene from the estimated *λ*′, *G*′. Inverting this typically requires the use of Bayes’ Theorem, but in this case we will assume that log *λ* and log *G* have a flat distribution and assume (from the Central Limit Theorem) that errors are small and normally distributed. In this case, we would like to determine the covariance matrix Σ′ for in terms of log *λ*′, log *G*′. We can calculate this taking a numerical derivative of Eq’s. S22 and S23. We estimate derivatives numerically, for instance *∂m/∂* log *λ* = (*m*(*λ* + *dλ, G*) *− m*(*λ, G*))*/d* log *λ*, typically sampled with *d* log *λ* = *dλ/λ* = 10^−6^. Then

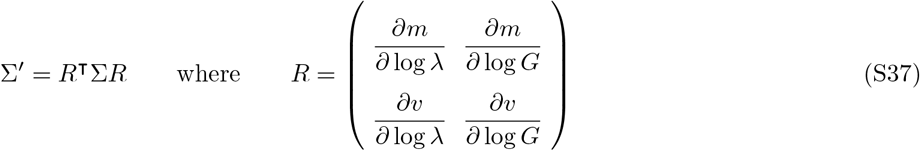

#### I. Assigning uncertainties

The covariance matrix and its inverse can be written in terms of the standard deviations *σ*_*x*_, *σ*_*y*_, and the Pearson correlation coefficient *ρ*.

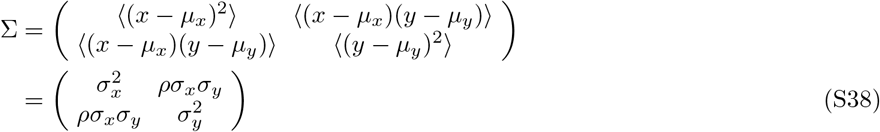

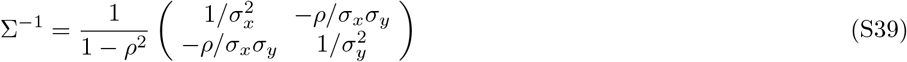

If we compare the possible values of *x* between two distributions with the same (possibly unknown) value of *y*, we need the standard deviation of the conditional distribution *P*(*x*, 0), which is 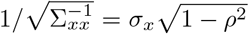. This would be the situation when comparing the change in gene expression for one gene between two samples with the same, possibly unknown, PCR bias. On the other hand, if we compare the possible values of *x* between two distributions with unkno<u>wn</u> and likely different values of *y*, we need the standard deviation of the marginal distribution *dy P* (*x, y*), which is 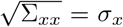. This would be the situation where we compare PCR gain between two samples with different expression levels.

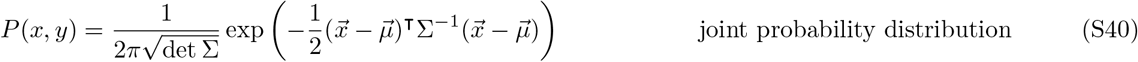

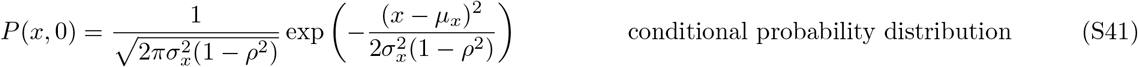

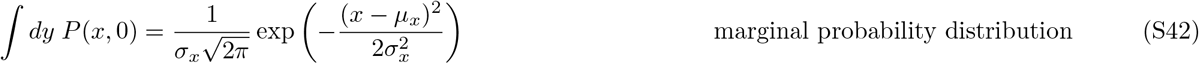

#### J. Limitations of the current approach

The previous section calculates the uncertainties of measuring *m*′, *v*′, *λ*′, *G* gives a known *λ* or *G*. In an experiment we only know the data and *m*′, *v*′ and seek a range of compatible values for *λ*′, *G*′. When the number of cells is large and standard errors are correspondingly small, we can plug the estimated values of *λ, G* into the real values in Eqs. S35 and S36. However, when the uncertainties are not small, when the number of cells is not great or when *λ ≪* 1 or *λ ≫* 1, this can cause substantial errors.

An extreme case is when we measure precisely zero reads among *N*_*cells*_ cells. The above analysis would erroneously estimate that both the mean and its uncertainty of *λ* were exactly zero (*m* = 0 and *σ*_*m*_ = 0). Yet such a measurement would be compatible with 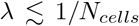, and therefore the correct uncertainty should be on the order of 1*/N*_*cells*_. Future work could use a Bayesian approach, though efforts were complicated by the excessive computational power required.

### II. PIPELINE

For each gene, the mean and variance of the log-normalized distribution of reads (Fig. S1b) serve as summary statistics, which are transformed into the estimates 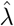 and 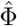 (Fig. S1c). The uncertainties *σ*_*λ*_ and *σ*_*NF*_ of 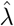 and 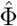, respectively, are estimated from the technical noise model (*σ*^model^) and from aggregating estimates between mice of the same age (*σ*^batch^), after adjusting for systematic changes in overall gene expression (Fig. S1d). Differential analysis of 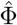 between different ages yields differential variance genes (DNGs), with significant fold changes typically ranging from 3 to 100 and 1/100 to 1/3 (Fig. S1e). The pipeline is tested against simulated gene expression calculated with technical noise but not biological noise, for artificial genes generated from the same distribution of *λ* and *NF* as the actual data (Fig. S1, bottom row). Each step of the pipeline is verified to be unbiased and properly calibrated within our technical noise model (Fig. 1a).

### III. ENRICHMENT ANALYSIS

DNGs are found in all tissues and cell types of *Tabula Muris Senis*, with a set of over a hundred universal DNGs frequently repeated across many cell types in the data set. We construct an enrichment analysis that takes into account the repetition of DNGs as well as the varying number of genes observed in each cell types. Our enrichment score counts the number of times that a DNG is in a given pathway, normalized by the number of genes that were tested for significance (potential DNGs).

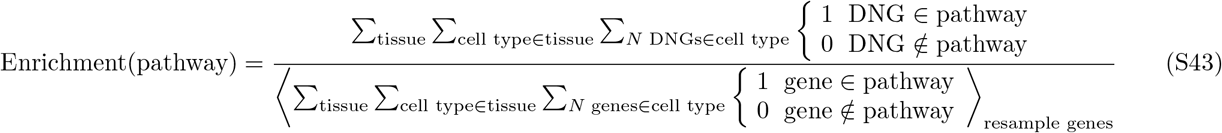

The numerator counts the number of times a DNG occurs in a given pathway, in each cell type, in each tissue. The denominator resamples genes with 10^−2^ *< λ <* 3 that were tested for differential variance, including actual DNGs. Resampling draws the same number of genes in each cell type and tissues as the number of DNGs, to ensure that the enrichment score averages to 1 for non-significant pathways. We further select only pathways for which at least 100 DNGs are counted with repitition, with at least 20 unique DNGs, and with a family-wise error rate less than 0.01 (Bonferroni correction), to minimize spurious results from overly narrow gene ontologies. Confidence intervals shown are not multiple hypothesis corrected, and enrichment scores are simulated with resampled data to ensure that the calculated FWER and confidence intervals are faithful. Pathways are downloaded from KEGG and Gene Ontology for mouse-specific genes and pathways.

### IV. ALTERNATIVE TO BENJAMINI–HOCHBERG PROCEDURE FOR MULTIPLE HYPOTHESIS TESTING

All statistical tests in this manuscript use the Benjamini–Hochberg procedure with false discovery rate *FDR* = *α* = 0.1, unless otherwise stated. The sole exception is choosing a number of high variance genes for clustering cell types. For this, we reconsider the notion that the *FDR* should be predetermined and fixed. In many situations in single cell analysis, small changes in the *FDR* can dramatically change the number of ‘significant’ genes, and thus strongly effects the sensitivity of the test. In other words, a small increase in *α* may cause many more true genes to be accepted than false ones. In these situations, it may be beneficial to relax the requirement of a fixed *FDR* if downstream analysis benefits from an increased sensitivity at the expense of a decreased specificity.

For selecting the number of genes for reclustering, we take *α* as a variable and calculate the number of significant genes *N*_*s*_(*α*). We estimate the true number of significant genes as the asymptote *N*_*s*_(*α →* 1) and estimate the true discovery rate *TDR* = *N*_*s*_(*α*)*/N*_*s*_(*α →* 1).

